# Structural basis of substrate transport and drug recognition by the human thiamine transporter SLC19A3

**DOI:** 10.1101/2024.03.11.584396

**Authors:** Florian Gabriel, Lea Spriestersbach, Antonia Fuhrmann, Katharina E. J. Jungnickel, Siavash Mostafavi, Els Pardon, Jan Steyaert, Christian Löw

**Affiliations:** European Molecular Biology Laboratory (EMBL) Hamburg, Notkestraße 85, 22607 Hamburg, Germany; Centre for Structural Systems Biology (CSSB), Notkestraße 85, 22607 Hamburg, Germany; Structural Biology Brussels, Vrije Universiteit Brussel (VUB), Brussels 1050, Belgium; VIB-VUB Center for Structural Biology, VIB, Brussels 1050, Belgium; Bernhard Nocht Institute for Tropical Medicine, 20359 Hamburg, Germany

## Abstract

Thiamine (vitamin B_1_) functions as an essential coenzyme in cells. Humans and other mammals cannot synthesise this vitamin *de novo* and thus have to take it up from their diet. Eventually, every cell needs to import thiamine across its plasma membrane which is mainly mediated by two specific thiamine transporters SLC19A2 and SLC19A3. Loss of function mutations in either of these transporters leads to detrimental, life-threatening metabolic disorders. SLC19A3 is furthermore a major site of drug interactions. Many medications, including antidepressants, antibiotics and chemotherapeutics are known to inhibit this transporter, with potentially fatal consequences for patients. Despite a thorough functional characterisation over the past two decades, the structural basis of its transport mechanism and drug interactions has remained elusive. Here, we report eight cryo-electron microscopy (cryo-EM) structures of the human thiamine transporter SLC19A3 in complex with various ligands. Conformation-specific nanobodies enabled us to capture different states of SLC19A3’s transport cycle, revealing the molecular details of thiamine recognition and transport. We identified nine novel drug interactions of SLC19A3 and determined structures of the transporter in complex with the inhibitors fedratinib, hydroxychloroquine, amprolium and amitriptyline. These data allow us to develop an understanding of the transport mechanism and ligand recognition of SLC19A3.

## Introduction

Thiamine, commonly known as vitamin B_1_, is crucial for the survival of cells and an essential micronutrient for humans and other mammals^1^. Inside the cell, thiamine is converted to thiamine pyrophosphate (thiamine-pp) by the enzyme thiamine pyrophosphokinase 1 (TPK1)^2,3^. Thiamine-pp then acts as a coenzyme in central metabolic pathways, such as the citric acid cycle and the pentose phosphate pathway^4^. Mammalian cells lack the ability to synthesise thiamine *de novo*. Instead, they import the vitamin over their plasma membrane through the transport activity of several integral membrane proteins of the solute carrier family (SLC, Fig. 1a)^5^. SLC19A2 and SLC19A3 are the main transporters for thiamine in humans and mediate the uptake of the vitamin with high affinity and specificity (K_m_ = 2-7 μM)^6–8^. They are closely related to the folate transporter SLC19A1, with which they form the SLC19A family of vitamin transporters^5^. In addition, the organic cation transporter 1 (OCT1, SLC22A1), which is known to accept a wide variety of substrates, can also mediate thiamine uptake, though with lower affinity (K_m_ ∼ 780 μM)^9^. Among the known thiamine transporters, SLC19A3 stands out in terms of its physiological and pharmacological importance, as it is crucial for the transport of thiamine across the intestinal wall and the blood-brain barrier^10,11^. Deletion of SLC19A3, but neither SLC19A2 nor OCT1, leads to strongly decreased thiamine levels in the blood serum and brain tissue under standard diet conditions^11–13^. Mutations of SLC19A3 cause severe neurometabolic disorders in the form of Wernicke’s-like encephalopathy (WLE)^14^ and biotin- and thiamine-responsive basal ganglia disease (BTBGD)^15–17^. In most cases, these diseases have an onset in infancy and can lead to life-long disabilities and early death^15^. Thiamine uptake through SLC19A3 is further known to be inhibited by a broad spectrum of commonly prescribed drugs, including antidepressants, antibiotics and antineoplastic medications^7,10^. Being under treatment with these thiamine uptake inhibitors (TUIs) can lead to drug-induced thiamine deficiencies on an organism-wide or tissue-specific level, with potentially fatal consequences^11^. This is exemplified by the case of the Janus-kinase (JAK) inhibitor fedratinib. Treatment with this compound induced the development of Wernicke’s encephalopathy, a severe neurodegenerative disorder characteristic for thiamine deficiency, in several patients^18^ (clinical trial ID: NCT01437787). This adverse event could eventually be linked to high-affinity inhibition of SLC19A3 by fedratinib^19^.

**Fig. 1:**
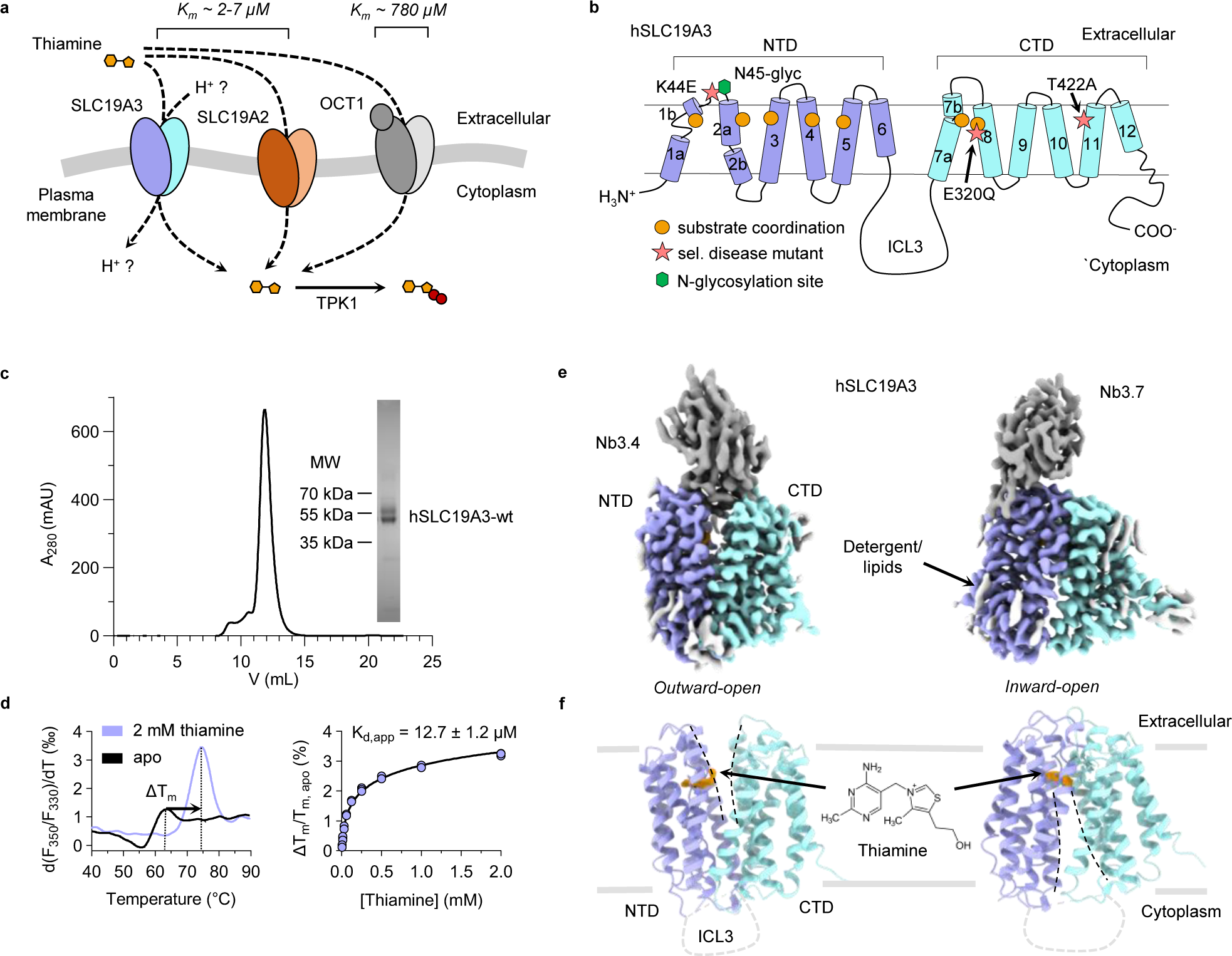
Structure and function of hSLC19A3. **a**, Humans and other mammals have at least three distinct membrane transporters that can mediate thiamine uptake: SLC19A3, SLC19A2 and OCT1 (SLC22A1). SLC19A3 is essential for the uptake of thiamine across the intestinal and blood-brain barrier under physiological conditions. In the cytoplasm, thiamine is phosphorylated by the enzyme TPK1 to form the biologically active coenzyme thiamine pyrophosphate. **b**, Cartoon representation of hSLC19A3. It follows the canonical MFS fold, with twelve transmembrane helices (TMs) folding in two symmetrically related six-helix bundle domains (NTD and CTD). Substrate coordination is mediated mostly by residues of TM1-5 of the NTD and by TM7 and TM8 of the CTD (orange circles). The positions of three disease mutants which were studied in this work are indicated by pink stars. **c**, Representative size-exclusion chromatography (SEC) trace of purified hSLC19A3. The protein elutes as a monomer of about 50 kDa. A fraction of the protein is glycosylated and appears as an extra band at ∼60 kDa in SDS-PAGE (inset). **d**, Thermal shift assay of hSLC19A3 using nanoDSF. The left panel shows the first derivative of the melting curve, measured as ratio of the fluorescence recorded at 350 nm (F350) and at 330 nm (F330), in the absence and presence of thiamine. Thiamine induces a strong stabilisation of the transporter (ΔT_m_ 10.9 ± 0.3 °C by 2 mM thiamine). The right panel illustrates the concentration dependent thermal shifts (*n = 3*) with the resulting apparent dissociation constant K_d,app_ for thiamine. **e**, Cryo-EM maps of thiamine-bound hSLC19A3 in its outward-open (left) and inward-open (right) conformation. The NTD (purple) and CTD (cyan) of the transporter, as well as the fiducial nanobodies (grey), are resolved. ChimeraX contour levels: outward-open: 0.876; inward-open: 0.148. **f**, Structure models of the respective conformational states. The density of thiamine is shown in orange. ChimeraX contour levels: outward-open: 0.547; inward-open: 0.148. The N-terminus, C-terminus and ICL3 are expected to be structurally disordered and could consequently not be resolved.

The human SLC19A3 (hSLC19A3) is a twelve-pass transmembrane protein of the Major Facilitator Superfamily (MFS)^20–22^. It is predicted to follow the canonical MFS fold, which consists of two α-helical and symmetrically related domains, termed the N-terminal (NTD) and the C-terminal domain (CTD), as shown in Fig. 1b. In recent years, the transport activity and drug-induced inhibition of SLC19A3 has been thoroughly studied on a cellular level^7,8,10,23–27^. The molecular basis of ligand recognition and the transport mechanism of SLC19A3 are, however, still not well understood.

Here, we report eight cryo-EM structures of hSLC19A3. Structure determination was enabled by the generation of three conformation-specific nanobodies against hSLC19A3. With these, we captured the solute carrier in its outward- and inward-open states, which provides unprecedented insights in its substrate recognition and transport cycle. Furthermore, we used thermal shift assays to screen for new drug interactions. This led to the discovery of nine novel high-affinity binders of hSLC19A3. To understand the structural basis of drug interactions of this transporter, we determined cryo-EM structures of hSLC19A3 in complex with the known high-affinity inhibitors hydroxychloroquine, fedratinib, amprolium and amitriptyline. These structural data allowed us to develop a first structure-based pharmacophore model of hSLC19A3.

## Results

### Cryo-EM structures of hSLC19A3 in different conformations

To study hSLC19A3 on a structural and biophysical level we made use of an expression system that has already been widely used for the functional characterisation of the transporter in cell-based assays^7,8,10,24^. Full-length hSLC19A3 was expressed in HEK293-derived Expi293F^TM^ cells and purified in a detergent solution using a mixture of lauryl maltose neopentyl glycol (LMNG) and cholesterol hemisuccinate (CHS) (Fig. 1c). The recombinant transporter is strongly stabilised against heat denaturation in the presence of its known substrate thiamine (ΔT_m_ = 10.9 ± 0.3 °C, Fig. 1d). Concentration-dependent thermal shift assays revealed high-affinity binding of thiamine with an apparent dissociation constant (K_d,app_) of 12.7 ± 1.2 μM (Fig. 1), which is in good agreement with previously reported transport affinities in the low micromolar range^8,9^. In contrast, phosphorylation of thiamine strongly reduced the interaction between the vitamin and hSLC19A3 (Supplementary Fig. 1). Since hSLC19A3 on its own proved to be too small for structure determination by cryo-EM (∼55 kDa), we generated hSLC19A3-specific nanobodies in a llama (Supplementary Fig. 2). For immunisation, a glycosylation-free mutant of hSLC19A3 (hSLC19A3-gf) was used in order to provide an increased accessible surface area for antibody binding (Supplementary Fig. 3). To generate hSLC19A3-gf, the predicted N-glycosylation sites Asn45 and Asn166 were mutated to glutamine residues. Thermal stability experiments did not show any interference of these mutations with thiamine binding (Supplementary Fig. 4). The immunisation and nanobody discovery were performed as described earlier^28^. In total we identified three hSLC19A3-specific binders: Nb3.3, Nb3.4 and Nb3.7 with binding affinities in the range of 100-300 nM (Supplementary Fig. 2). While Nb3.3 and Nb3.4 bind stably to wildtype hSLC19A3, Nb3.7 only interacts with the glycosylation-free transporter (binding data not shown). All three nanobodies were used for structure determination of hSLC19A3 by single particle cryo-EM with map resolutions reaching between 2.9-3.8 Å on a global level and 2.4-3.2 Å in the substrate binding site (Supplementary Fig. 5-13, Supplementary table 1). Nb3.4 stabilises the outward-open state of the transporter, while Nb3.3 and Nb3.7 both bind to its inward-open state (Fig. 1e,f, Supplementary Fig. 6-9). Using this molecular toolset, we determined the structures of hSLC19A3 in its outward-open and inward-open conformations, in both apo and thiamine-bound states.

The experimental structures of hSLC19A3 confirm the MFS fold of the transporter, with twelve transmembrane helices (TM) folding into two compact six-helix bundle domains (Fig. 1f, Supplementary Fig. 14). The domains are connected via a 76-residues long (Lys195-Glu271), poorly conserved intracellular loop 3 (ICL3) (Supplementary Fig. 15). No density could be resolved for this loop, indicating that this part of the transporter is likely disordered and flexible. Similarly, parts of the N-terminus (Met1-Ser10) and the C-terminus (Tyr459-Leu496) could not be resolved in any of the cryo-EM structures (Fig. 1e,f). Within the well resolved helical domains, there are three discontinuous transmembrane helices in both the outward- and inward-open state: TM1, TM2, and TM7. In TM1, Ile13-Met27 form a stable α-helix (TM1a). The helix then partially unwinds and is followed by a short helical segment (TM1b), which is flanked by two proline residues in position 33 and 42 (Supplementary Fig. 14a). TM2 is interrupted close to its cytoplasmic end, between Val65 and Leu68. This discontinuity allows TM2 to bend around TM4 (Supplementary Fig. 14a). The helical structure of TM7 is also broken, in this case close to the extracellular space, between Asn297 and Gln300 (Supplementary Fig. 14b). Our structural data further confirm that recombinant hSLC19A3 is N-glycosylated on Asn45, whereas the second predicted glycosylation side, Asn166, appears to be glycosylation-free in the wildtype transporter (Supplementary Fig. 3). Additional non-protein density observed in the EM-reconstructions likely originates from bound lipids or detergent molecules (Fig. 1e).

### Thiamine binding site

Comparison of the final cryo-EM reconstructions from data sets recorded in the absence and presence of thiamine revealed extra densities in the transporter vestibule which only appeared in the thiamine-supplied samples. These densities correspond well to the molecular shape of thiamine. The fitted ligand also matches with the transporter structures in terms of coordinating interactions (Fig. 2). These structures highlight that the substrate binding site is positioned close to the extracellular side of hSLC19A3 (Fig. 2a,b). Thus, the thiamine binding site in hSLC19A3 is similar to the reported substrate binding pocket of the closely related folate transporter SLC19A1^29–31^ (Supplementary Fig. 16b,d,f). This is, however, a rather unusual feature for MFS transporters, which typically bind their substrates more centrally within the membrane plane^32^ (Supplementary Fig. 17). Since the discovered nanobodies allowed the determination of different conformational states of hSLC19A3, a direct comparison of thiamine binding in the outward-open and the inward-open state was possible.

**Fig. 2:**
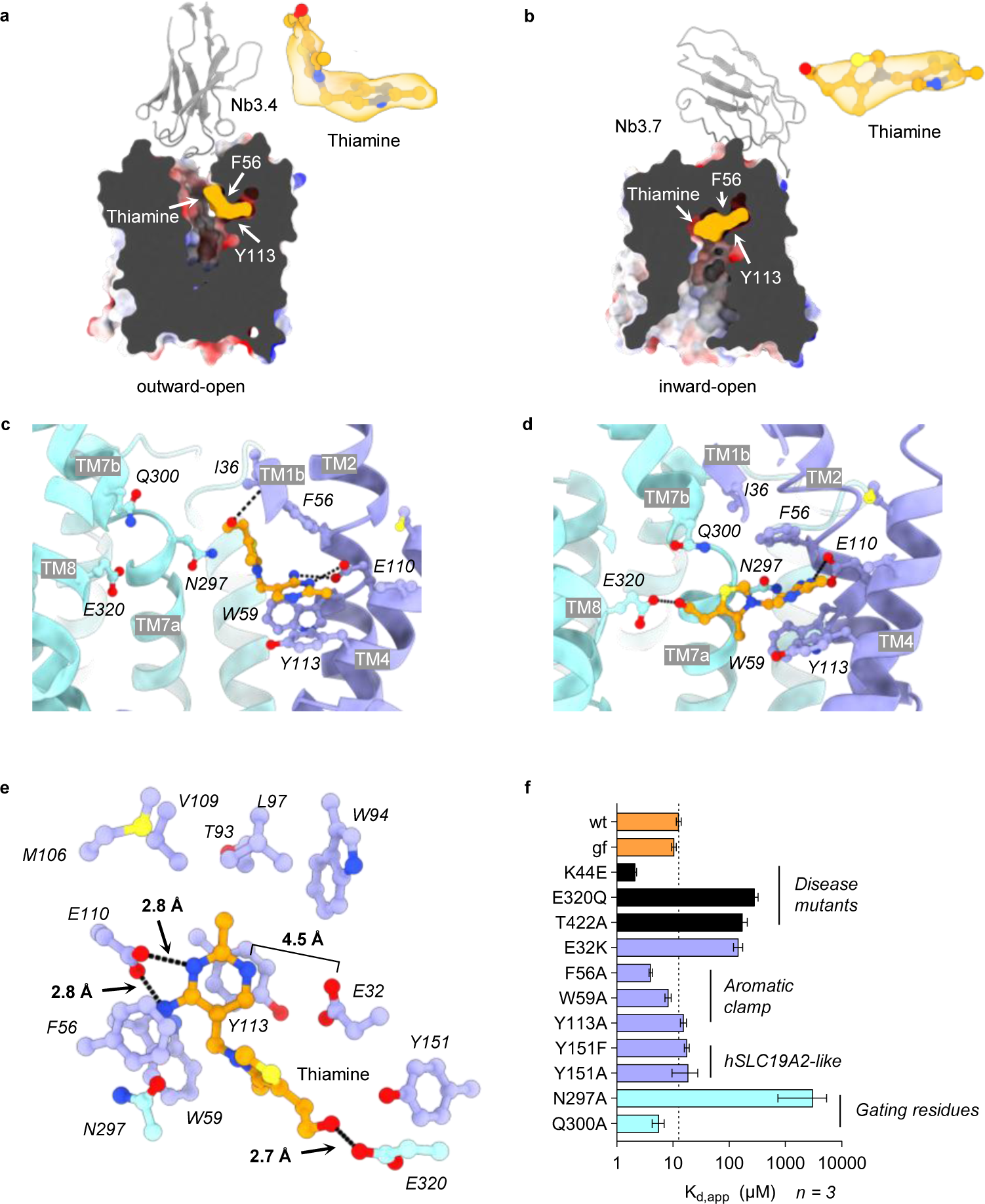
Thiamine binding site of hSLC19A3. **a**, Localisation of the thiamine binding site of hSLC19A3 in the outward- and **b**, inward-open conformation of the transporter. Density for thiamine (orange) is shown separately (ChimeraX contour levels: 0.547 and 0.148). **c**, Side view of residues involved in thiamine coordination in the outward- and **d**, inward-open conformation of hLSC19A3. **e**, Top view of hSLC19A3 highlighting the thiamine-coordinating residues in the substrate binding site of the transporter. Residues of the NTD are coloured in purple, residues of the CTD in cyan, and thiamine in orange. Black dashes indicate hydrogen bonds (cut-off at 3.0 Å)^33^ **f**, Thiamine binding affinities (apparent dissociation constants K_d,app_) of single point mutants of hSLC19A3 (*n = 3*, independent measurements, ± s.d.).

In the outward-open state, thiamine is exclusively coordinated by the N-terminal domain (NTD) of the transporter (Fig. 2a,c). Thiamine adapts a kinked conformation with regard to its two aromatic rings. Its aminopyrimidine moiety inserts into a complementary pocket of the NTD, parallel to the membrane plane (Fig. 2a,c). Here, the aromatic ring interacts via π-π stacking with Tyr113 and to a lesser extend with Trp59 (Fig. 2c,e). The methyl group of the aminopyrimidine ring extends into a hydrophobic pocket formed by the side chains of Thr93, Trp94 and Leu97 on TM3, and Met106 and Val109 on TM4 (Fig. 2e). Crucial polar contacts are formed with Glu110, which is within hydrogen bonding distance (≤ 3.0 Å)^33^ to both the ring-nitrogen (2.8 Å), as well as the primary amine of thiamine (2.8 Å) (Fig 2e). Based on the resolved structures and the observed interaction with thiamine, it is likely that Glu110 is protonated. This is supported by predicted neutral pK_a_ values of Glu110 (pK_a_ = 6.4-7.5) throughout the transport cycle (structure-based pK_a_ estimation using PROPKA^34^, see Supplementary Fig. 18). The hydrogen bonding between the aminopyrimidine ring and Glu110 appears to be critical and specific for the high-affinity binding of thiamine to hSLC19A3. In oxythiamine, a close derivative of thiamine, the primary amine of the aminopyrimidine ring is substituted by a carbonyl oxygen. This subtle change already leads to a strongly decreased affinity and a lower thermostabilisation of the transporter (Supplementary Fig. 1b,e). Opposite of Glu110, Glu32 provides another polar, solvent exposed surface in the substrate binding site of hSLC19A3. However, thiamine does not provide any strong hydrogen bond donors for pairing with this residue and is also out of hydrogen bond distance (Fig. 2e). In the outward-open state of hSLC19A3, the thiazolium ring of thiamine is oriented approximately 90° relative to the aminopyrimidine ring and is primarily coordinated through π-π stacking with Phe56. The hydroxyethyl group, the tail of thiamine, reaches even further up in the binding site, forming polar contacts with the backbone of Ile36 (Fig. 2c).

When hSLC19A3 transitions from the outward- to the inward-open state, thiamine undergoes a major rearrangement. It relaxes via its rotatable *C-C* and *C-N* bonds connecting the aminopyrimidine and thiazolium ring into an elongated conformation (Fig. 2d,e). The aminopyrimidine ring stays coordinated in the same position in the NTD, with Phe56 closing on top of it. Together with Trp59 and Tyr113, Phe56 forms now a hydrophobic pocket that we termed ‘aromatic clamp’ (Fig. 2e). Meanwhile, the positively charged thiazolium ring reorients and is accommodated between the electronegative backbone and side chain carbonyl oxygen atoms of Leu296 and Asn297, respectively. Through the transition of hSLC19A3 to the inward-open conformation, the hydroxyethyl tail of thiamine detaches from the backbone of Ile36 and reaches across to Glu320 on the CTD to form a hydrogen bond (2.7 Å). With this, thiamine tightly connects the two helical bundles of the transporter by non-covalent interactions (Fig. 2d,e). Glu320 is further predicted to undergo strong pK_a_ changes over the course of the transport cycle, which could link thiamine transport to a proton gradient over the membrane (see discussion and Supplementary Fig. 18).

In order to get experimental insights into the interaction of individual residues with thiamine, we generated and analysed several single point mutants of hSLC19A3 (Fig. 2f, Supplementary Fig. 4 and 19). The mutation of residues of the aromatic clamp to alanine did not impair thiamine binding (Fig. 2f). We speculate that two out of its three constituents, Phe56, Trp59 and Tyr113, are sufficient to mediate stable π-π stacking to the aminopyrimidine ring of thiamine. However, mutating Asn297 to alanine, and thereby removing a negative partial charge from the substrate binding site, led to a ∼ 250-fold loss in binding affinity (Fig. 2f). In addition, we probed the role of Glu32 in thiamine binding. The mutation of the equivalent residue Glu45 to lysine in the closely related folate transporter SLC19A1 leads to a loss of function and conveys methotrexate (MTX) resistance^35,36^ (Supplementary Fig., 16). The analogous mutation (hSLC19A3-Glu32Lys) causes a 10-fold drop in the affinity for thiamine in hSLC19A3, from 12.7 ± 1.2 μM to 145 ± 27 μM. It is likely that the placement of a positive charge in this position leads to an electrostatic or partially steric repulsion of the aminopyrimidine ring (Fig. 2e,f). The interaction between Glu320 and the hydroxyethyl tail of thiamine is affected in a rare form of heritable Wernicke’s-like encephalopathy^14^. Our biophysical data illustrate that the disease associated mutation Glu320Gln leads to a 20-fold decrease in affinity for thiamine, from 12.7 ± 1.2 μM to 284 ± 45 μM (Fig. 2f).

### Substrate binding sites in other human thiamine transporters

The transport of thiamine can be mediated by different transporters, dependent on the tissue, cell type and subcellular compartment^23,25,37,38^ (Fig. 1a). High-affinity transport of thiamine across the plasma membrane is shared between SLC19A3 and the closely related SLC19A2^6,26^. In order to understand the structural relationship between these two, we compared the experimentally determined structure of the inward-open hSLC19A3 with the AlphaFold2 (AF2) predicted structure of hSLC19A2^39^ (Supplementary Fig. 16 and 20).

The global fold of the two proteins is almost identical, with the only marked differences in the poorly conserved and disordered N- and C-terminal regions, as well as ICL3 (Supplementary Fig. 16b). The substrate binding site appears to be structurally highly conserved. Only subtle differences in the form of two substitutions, (i) Phe56^hSLC19A3^ to Tyr74^hSLC19A2^ and (ii) Tyr151^hSLC19A3^ to Phe169^hSLC19A2^ can be observed (Supplementary Fig. 16e). The first substitution introduces a minor change in the aromatic clamp. As shown in Fig. 2f, replacing Phe56^hSLC19A3^ with alanine in hSLC19A3 did not impact the binding affinity for thiamine. Therefore, we assume that the even more similar substitution of Phe56 to tyrosine will not impact substrate binding. The effect of the second substitution (ii) on thiamine binding was tested experimentally. When Tyr151^hSLC19A3^ was replaced with phenylalanine, no strong effect on the affinity for thiamine was observed (Fig. 2f). This in line with previous observations, which reported very similar transport affinities for the two thiamine transporters^7,8^. The situation is different, when comparing the hSLC19A3 structures with the recently determined cryo-EM structure of OCT1 (SLC22A1) in complex with thiamine. The reported transport affinity of OCT1 for thiamine is much lower than for the SLC19A transporters (K_m_ ∼ 780 μM compared to 2-7 μM)^7–9^. This difference could be explained by a looser coordination of thiamine in OCT1, compared to the tight binding observed in hSLC19A3^40^ (Supplementary Fig. 17).

### Transport cycle

To obtain a better understanding of the conformational changes during the transport cycle we compared the cryo-EM structures of hSLC19A3 stabilised in the outward-open state by Nb3.4 with the inward-open state, bound by Nb3.3 or Nb3.7. An overlay of the individual N- and C-terminal domains of the different conformational states revealed no major intradomain rearrangements (C_α_ RMSD of 0.52-1.94 Å). Only extracellular loop 4 (ECL4) and intracellular loop 5 (ICL5) reorient slightly (Supplementary Fig. 14). Based on the available structures, we conclude that the conformational changes observed for hSLC19A3 follow a rocker-switch mechanism, in which the NTD and CTD move as rigid bodies to create a moving barrier around the substrate binding site^32^. This barrier consists of a cytoplasmic gate and an extracellular gate that open and close in an alternating fashion. This provides alternating solvent accessibility to the substrate binding site from the extracellular and cytoplasmic space (Supplementary Fig. 21). The cytoplasmic gate of hSLC19A3 is formed by hydrophobic contacts of Tyr128 (TM4) and Ala395 (TM10) with Tyr403 (TM11). The gate is further stabilised by polar interactions of the side chains of Asp75 (TM2) and Gln137 (TM5) with the backbone of Ala404 and Leu405 (TM11) and Lys338 (ICL5), respectively (Supplementary Fig. 21a-c). These contacts are broken when the transporter transitions to the inward-open state (Supplementary Fig. 21d-f). The long ICL3 of hSLC19A3 allows for a dilation of the cytoplasmic gate. Simultaneously, a barrier to the extracellular space is established by the closure of the extracellular gate. The core of this gate is built by polar contacts of the side chains of Asn297 and Gln300 (TM7). They are within hydrogen bonding distance with the backbone of Phe56 (TM2) and Pro33 (TM1a), respectively. These polar contacts are supported by hydrophobic interactions between Ile36 (TM1b) and Ile301 (TM7). With these in place, TM1b is coordinated as a lid, sealing the extracellular gate (Supplementary Fig. 21d-f).

### Mutation of hSLC19A3 in biotin- and thiamine responsive basal ganglia disease (BTBGD)

Loss of function mutations of hSLC19A3 cause BTBGD in humans^15,17,41–45^ (Supplementary Fig. 22). This disease is marked by severe neurological symptoms and is inherited genetically in an autosomal recessive manner^15^. The median age of onset of symptoms varies between three months and four years after birth, dependent on the underlying mutations in the SLC19A3 gene. Under the currently advised treatment - a combination therapy of high-dose thiamine and biotin - more than 30% of the patients still suffer from moderate to severe effects of the disease, while about 5% of those affected die despite treatment^15^. To gain a better understanding of the molecular cause of this disorder, we investigated the effects of one of the most common BTBGD-causing point mutation SLC19A3-Thr422Ala. Most cases are reported in Saudi Arabia, where the heterozygous carrier frequency is about 1 in 500 among the Saudi population^17^. The patients usually present with symptoms including subacute encephalopathy with confusion, convulsions, dysarthria and dystonia^46^. On a cellular level, the SLC19A3-Thr422Ala mutant localises normally to the (apical) plasma membrane in Caco-2 and MDCK cells^25^ and can also be expressed and purified *in vitro* (Supplementary Fig. 19a). However, its thiamine uptake activity is significantly impaired^25^. This agrees with our data, which highlight that the Thr422Ala mutation leads to a > 10-fold decrease of binding affinity for thiamine, from 12.7 ± 1.2 μM to 172 ± 35 μM (Fig. 2f, Supplementary Fig. 23d). Thr422 is located on TM11 and thus spatially in the outer region of the transporter. It is, however, physically connected to the core of hSLC19A3 by hydrogen bonding with the side chain of Gln294 on TM7 (Supplementary Fig. 22b,c and 23d). This part of TM7 is involved in the formation of the substrate binding site and the extracellular gate through Asn297, Gln300 and Ile301 (Fig. 2c,d, Supplementary Fig. 21). Structural changes in this region might explain the loss of affinity and transport activity of the Thr422Ala mutant.

The cryo-EM structures determined in this study additionally provide first hints on the disease-causing mechanism of other BTBGD mutations. Trp59Arg, Trp94Arg, Tyr113His, and Glu320Lys would be directly affecting thiamine binding by placing a positive charge in the substrate binding pocket of hSLC19A3 (Fig. 2e, Supplementary Fig. 22b,c). Binding data on the previously mentioned Glu32Lys mutant illustrate that the insertion of a positive charge in this region can lead to a strong loss of affinity for thiamine (>10-fold, Fig. 2f, Supplementary Fig. 4). This would likely render the transporter unreceptive for physiological thiamine concentrations, which is the possible cause for the pathogenicity of theses mutations.

### Drug interactions of SLC19A3

Previous studies identified a broad spectrum of FDA-approved drugs that act as thiamine uptake inhibitors (TUIs) by blocking hSLC19A3^7,10,11^. This has important clinical implications, as many of these drugs are widely prescribed over extended periods of time. Among these drugs are antidepressants, such as sertraline (Zoloft® and others) and amitriptyline, the antiparasitic hydroxychloroquine and the Janus-kinase (JAK) inhibitor fedratinib (Inrebic®). The inhibition of SLC19A3 can lead to organism-wide and tissue-specific thiamine deficiencies. In the latter case, the deficiencies would not be reflected in serum thiamine levels and remain undetected in standard diagnostics. As BTBGD-causing mutations and the example of SLC19A3 knock-out mice illustrate, this can lead to the decline of entire cell populations in vital organs, particularly in the brain^11,47^. Studying recombinantly purified hSLC19A3 in thermal shift assays, we explored the interaction space of the transporter with six known TUIs and eleven pharmacologically related drugs (Fig. 3). The corresponding screen confirmed physical binding of the known inhibitors to hSLC19A3, and further led to the identification of nine novel drug interactions with potential inhibitor function (Fig. 3a). To begin with, several drugs that are structurally related to thiamine were probed. The known hSLC19A3 inhibitors amprolium and trimethoprim strongly stabilised the transporter against heat unfolding in a concentration-dependent manner. This effect was exceeded by the newly tested antiparasitic pyrimethamine (Fig. 3a). Even at a concentration of 20 μM, this compound led to a thermostabilisation of 9.3 ± 0.5 °C and is thus a new potential inhibitor of hSLC19A3. Next, representative JAK inhibitors were analysed. Fedratinib, which is known to be a potent inhibitor of hSLC19A3, also showed high-affinity binding to the transporter in the thermal shift assays (K_d,app_ = 1.0 ± 0.2 μM). The interaction of SLC19A3 with three other JAK inhibitors, tofacitinib (Xeljanz®), baricitinib (Olumiant®) and momelotinib (Ojjaara®) was tested as well. These compounds are functionally similar to, but structurally distinct from fedratinib. Tofacitinib is also stabilising hSLC19A3, but only at higher concentrations (Fig. 3a). Baricitinib and momelotinib did not stabilise hSLC19A3 and thus likely do not interact with the transporter. This agrees with previous cellular uptake inhibition studies of JAK inhibitors^7^.

**Fig. 3:**
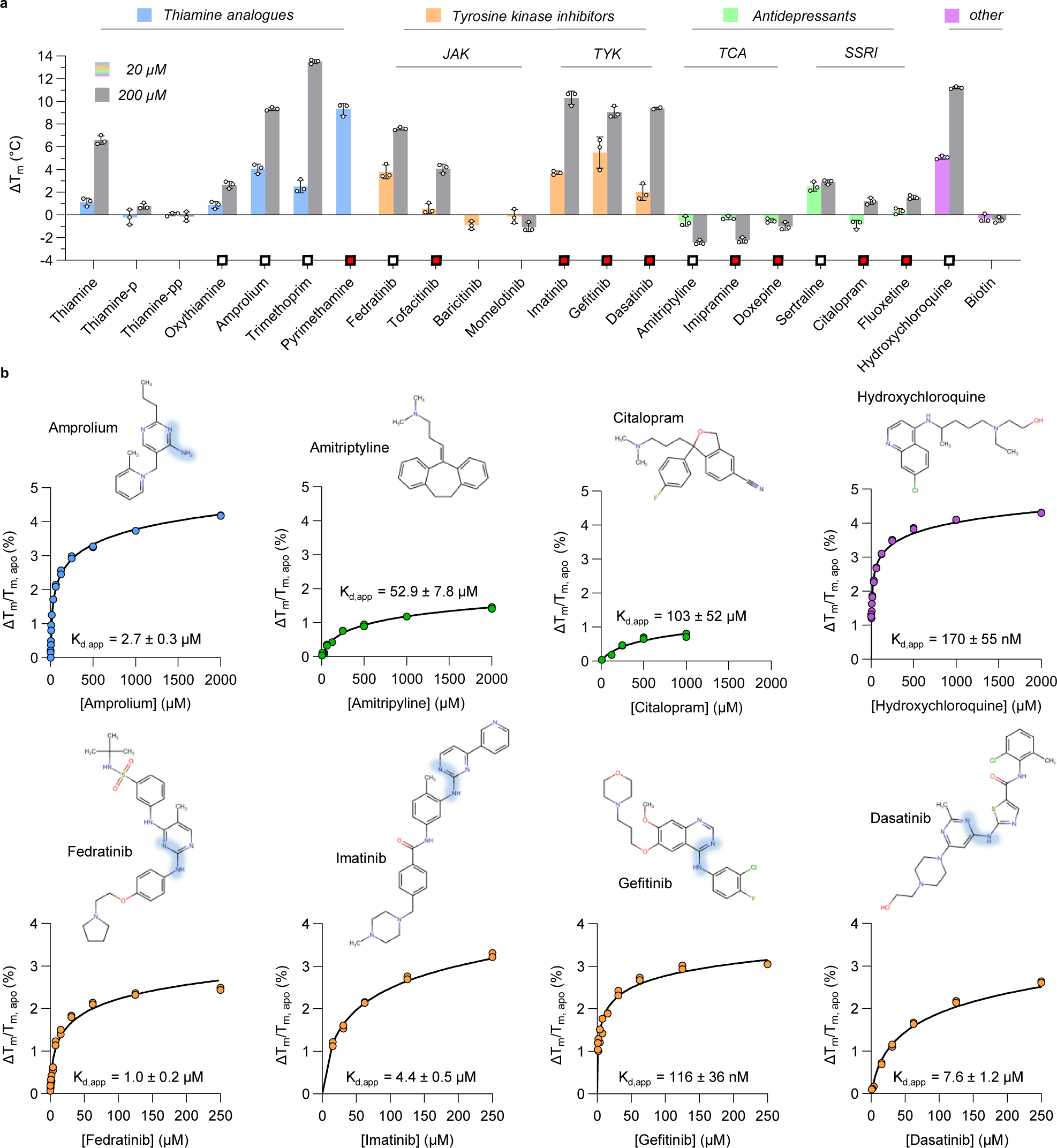
Drug interactions of hSLC19A3. **a**, Thermal shift screen of known and potential thiamine uptake inhibitors (TUIs). Thermal shift assays were performed in the presence of 20 μM and 200 μM of the respective compound and the stabilisation effect compared to the apo stability of hSLC19A3. The melting temperature was determined as the inflection point of the thermal unfolding curve of the protein (cf. Fig. 1d). Wherever possible, the ratio of the fluorescence at 350 nm and 330 nm (F350/F330) was selected as a readout for the thermal unfolding curve. Due to intrinsic fluorescence of the following compounds, the fluorescence trace at 350 nm (F350) only was used as a readout: oxythiamine, trimethoprim, pyrimethamine, tofacitinib, imatinib, gefitinib, dasatinib, sertraline, fluoxetine and hydroxychloroquine. The two compounds, pyrimethamine and baricitinib, interfere at 200 μM too strongly with the fluorescence detection to provide an unambiguous readout and were omitted in this dataset. The squares indicate identified TUIs. White squares represent already known TUIs, and red squares newly identified ones. **b**, Determination of the apparent binding affinity (K_d,app_) of the TUIs, identified in the thermal shift screen, using the readouts as described above. Shown are the chemical structures and the binding isotherms that were fitted according to a modified script of Hall’s method (Hall, 2019, Kotov et al., 2023) to the experimentally determined compound concentration-dependent thermal shifts (*n = 3*, independent measurements, shown as coloured circles). The blue highlights in the chemical structures of amprolium, fedratinib, imatinib, gefitinib and dasatinib indicate the presence of an amine adjacent to an unprotonated ring nitrogen, a shared chemical feature of many TUIs^7^.

Another important drug class that was tested were tyrosine kinase inhibitors, represented by imatinib (Gleevec®), gefitinib (Iressa®) and dasatinib (Sprycel®). They strongly stabilized hSLC19A3 with binding affinities of 4.4 ± 0.5 μM, 116 ± 36 nM and 7.5 ± 1.5 μM, respectively (Fig. 3b). These affinities are similar to the structurally related fedratinib. When comparing fedratinib and the tyrosine kinase inhibitors on a chemical level, they share a central aminopyrimidine moiety (highlighted in blue in Fig. 3b). This moiety is also at the core of the interaction of thiamine within the substrate binding site (Fig. 2e) and is thus potentially important for the interaction of the tyrosine kinase inhibitors with SLC19A3, as already suggested by Giacomini and co-workers (Giacomini et al, 2017).

All of the tested tricyclic antidepressants (TCA), including the known inhibitor amitriptyline and the newly assessed ones, imipramine and doxepin, interact with hSLC19A3 (Fig. 3a). Interestingly, TCAs actually destabilize the transporter in a concentration-dependent manner, while compounds of other classes were generally stabilising. In the group of selective serotonin reuptake inhibitors (SSRI), sertraline stabilises the hSLC19A3 already at concentrations in the low micromolar range (ΔT_m_ = 2.5 ± 0.4 °C), which is in agreement with existing data on cellular thiamine uptake inhibition^10^. The newly tested citalopram (Celexa® and others) and fluoxetine (Prozac® and others) also interact with hSLC19A3, but only at higher concentrations (Fig. 3a). The last analysed TUI was hydroxychloroquine (Fig. 3b). This drug strongly stabilises hSLC19A3 and binds to the transporter with high-affinity in the nanomolar range (K_d,app_ = 170 ± 55 nM). This agrees with the previously reported high-affinity inhibition of hSLC19A3 by hydroxychloroquine^10^. In contrast to the known and newly identified hSLC19A3 inhibitors, biotin does not interact with the transporter (Fig. 3b), which is in agreement with existing cellular uptake data^25^.

### Structural basis for thiamine uptake inhibition

The identified thiamine uptake inhibitors (TUIs) are structurally diverse (Fig. 3b). Their chemical heterogeneity presents a challenge for the development of a universal ligand-based pharmacophore model^7^. To elucidate how these compounds bind to hSLC19A3, we determined cryo-EM structures of the transporter in complex with four known transport inhibitors: (i) hydroxychloroquine, (ii) fedratinib, (iii) amprolium and (iv) amitriptyline (Fig. 4a-d). They represent structurally and functionally different drug classes. Cryo-EM structures were solved using glycosylation-free hSLC19A3-gf (see methods) in complex with Nb3.7, which stabilises the transporter in the inward-open state. The cryo-EM maps of the transporter-drug complexes were resolved to 3.1-3.7 Å, reaching resolutions between 2.5-3.2 Å in the core of the protein (Supplementary Fig. 10-13). All of the analysed inhibitors bind orthosterically in the substrate binding site of the transporter (Fig. 4). From a structural point of view, the binding of these compounds to hSLC19A3 is determined by at least three key factors, which are summarised in Fig. 4e,f.

1. Intercalation of aromatic rings in the aromatic clamp. Our cryo-EM data show that the aromatic rings of all four structurally resolved inhibitors form direct π-π stacking interactions with Tyr113 and to a lesser extent with Phe56 and Trp59 (Fig. 4a-d). This seems to be a universally important feature of the interaction of TUIs with hSLC19A3, as all known and newly identified ligands comprise at least one aromatic ring. This interaction seems particularly important for the binding of the transporter to compounds that otherwise lack compatible polar contacts. This concerns specifically the tested antidepressants amitriptyline, imipramine, doxepin, sertraline, citalopram, and fluoxetine (Fig. 3a)
2. Electrostatic compatibility with the polar contact points provided by Glu32, Glu110 and Asn297. Hydroxychloroquine, fedratinib and amprolium provide hydrogen bond donors and acceptors that pair spatially well with the polar residues of the substrate binding site (Fig. 4a-c). All three inhibitors are within hydrogen bonding distance to Glu110. As mentioned above, this residue is likely protonated throughout the transport cycle of hSLC19A3 (Supplementary Fig. 18) allowing Glu110 to act simultaneously as a hydrogen bond donor and acceptor. This agrees with the observed interaction patterns of fedratinib and amprolium with this residue (Fig. 4a). Another important polar residue of the substrate binding site is Asn297. Its side chain carbonyl oxygen contributes to the electrostatically negative interaction surface of the transporter in general, and with 2.8 Å, it is within hydrogen bonding distance with the primary amine of amprolium (Fig. 4c). The coordination capability of Glu320 is essential for high-affinity binding of thiamine, as shown in Fig. 2f and Supplementary Fig. 23c. For inhibitor binding, however, this residue seems to play no major role, as no interactions between its side chain and the inhibitors could be observed (Fig. 4a-d). Glu32, in contrast, is in clear hydrogen bond distance with one of the secondary amines of fedratinib (2.8 Å, Fig. 4b).
3. Insertion of a lipophilic moiety in the hydrophobic pocket formed by Thr93, Trp94, Leu97, Met106 and Val109. Three of the structurally resolved inhibitors engage in this type of interaction and protrude an apolar substituent into the hydrophobic pocket (Fig. 4a-c), namely a chlorine (hydroxychloroquine), a methyl (fedratinib) and a propyl group (amprolium). This interaction is likely beneficial for high affinity binding to hSLC19A3. A structurally corresponding methyl group can, for example, also be found in the chemical structures of the high-affinity binders imatinib and dasatinib (Fig. 3b).

**Fig. 4:**
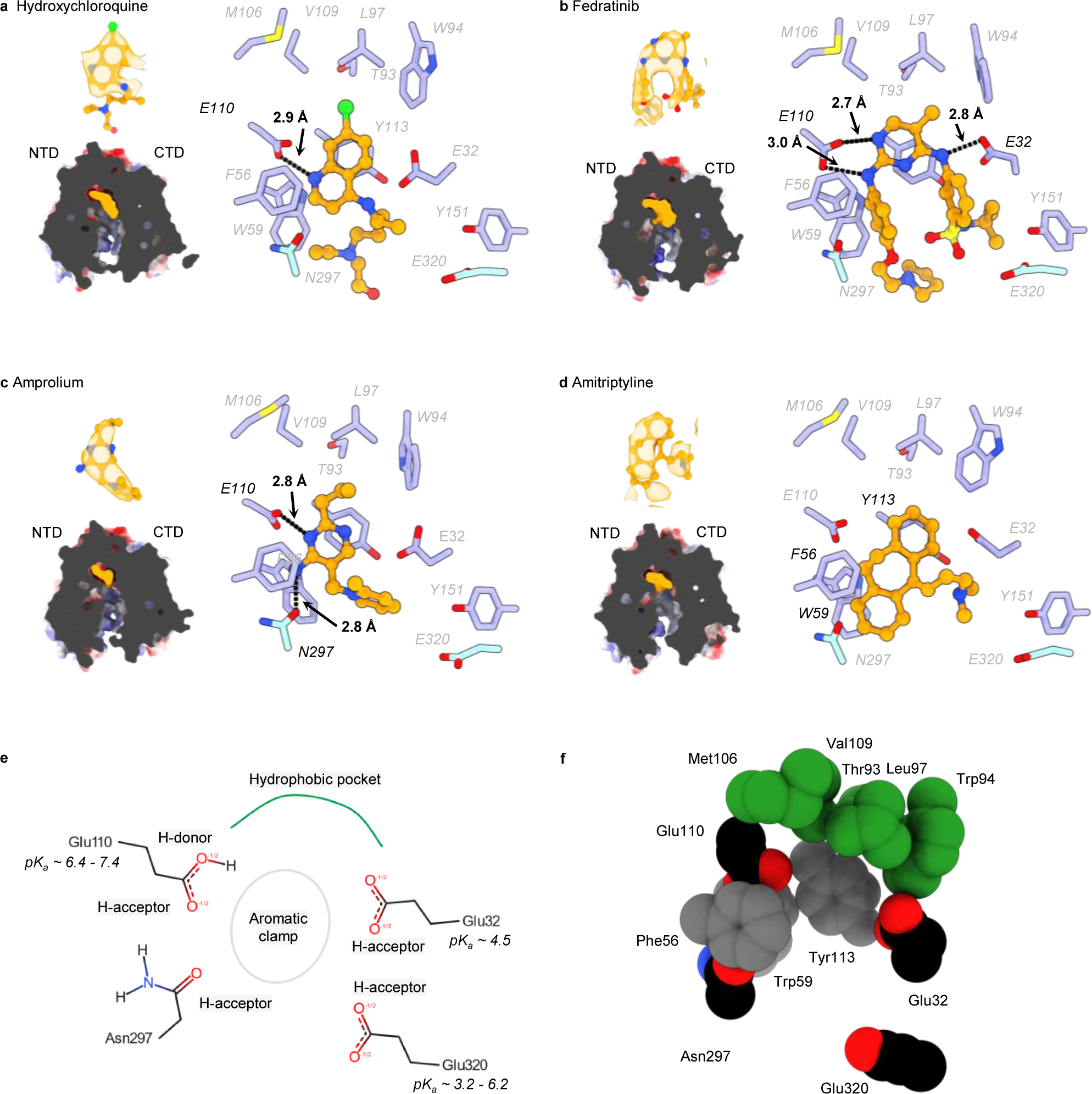
Cryo-EM structures of hSLC19A3 in complex with high-affinity inhibitors. **a-d,** Bound inhibitors are illustrated (orange) and were observed to occupy the same substrate binding site and engage with similar residues as thiamine. Shown are cross sections of the electrostatic surface of the ligand-bound structures (*left*), with the ligand coloured in orange; the densities for the different compounds are shown in the centre of the respective panels (ChimeraX contour levels for hydroxychloroquine: 0.128, fedratinib: 0.604, amprolium: 0.1, amitriptyline: 0.767); the coordination of the compounds is highlighted in the substrate binding site. Residues of the NTD are shown in purple, the ones of the CTD in cyan and the compound in orange. Black dashes indicate hydrogen bonds (cut-off at 3.0 Å)^33^. **e**, Structure-based pharmacophore model of hSLC19A3. Different molecular modules important for substrate and drug interactions of hSLC19A3 can be identified: the aromatic clamp, a hydrophobic pocket, a hydrogen bond donor (H-donor) and four hydrogen bond acceptors (H-acceptors). For the three involved glutamate residues, pK_a_ values were estimated for all resolved structures using PROPKA (Bas et al., 2008). In all structures, Glu32 and Glu110 have a constant pK_a_ of 4.5 and 7.4, respectively. In contrast, Glu320, exhibits a strong pK_a_ shift from 6.2 in the outward-open conformation to 3.2 in the inward-open state of hSLC19A3 (Supplementary Fig. 18). **f**, 3D structure representation of the key residues involved in drug recognition by hSLC19A3. Highlighted are the residues of the aromatic clamp (grey), the hydrophobic pocket (green) and the polar contact points of the substrate binding site (coloured by element, carbon: black, nitrogen: blue, oxygen: red).

## Discussion

Our structural and biophysical data reveal key aspects of the transport cycle and drug interactions of hSLC19A3. The transporter moves in a rocker-switch motion over the course of its transport cycle, creating a moving barrier around its substrate (Fig. 2, Supplementary Fig. 14, 18 and 21). Our data support a model where thiamine is first recognised via the NTD in the outward-open hSLC19A3. Once the transporter changes to its inward-open conformation, thiamine rearranges to form a molecular bridge between the NTD and the CTD, before it dissociates into the cytoplasm. The binding of thiamine follows a key-in-lock mechanism, with almost no structural differences between apo and substrate-bound structures of the transporter (C_α_ RMSD of 0.30-0.62 Å; Supplementary Fig. 14). We identified the key residues mediating thiamine recognition in hSLC19A3. Several of these are mutated in severe neurometabolic diseases (BTBGD and WLE, Supplementary Fig. 22). Regarding the transport mode of hSLC19A3, it is still not clear whether the transporter operates in a uniporter fashion or requires co-substrates to drive thiamine uptake. Energetically, a uniporter mechanism is plausible. The membrane potential in combination with the concentration gradient of thiamine might be sufficient for uptake of this organic cation across the plasma membrane. Our data highlight that thiamine is not able to bind to hSLC19A3 once it is phosphorylated (Supplementary Fig. 1). With this, the exit route for the vitamin via hSLC19A3 is blocked and shifts the transport equilibrium in favour of thiamine import (Fig. 1a).

A coupling of the transport mechanism of hSLC19A3 to sodium ions was not observed in a previous study^25^. There are conflicting data about the role of protons for the activity of hSLC19A3. While some works indicate both proton-dependent anti- and symport mechanisms for the transporter^8,48^, others see no link between the uptake of thiamine and proton gradients^24^. Based on the structural work presented here, we provide evidence that the binding site residue Glu320 undergoes a strong pK_a_ shift (from 6.2 to 3.2) over the course of the transport cycle and thereby potentially connects the uptake of thiamine to the symport of protons across the membrane (Supplementary Fig. 18). However, functional studies, ideally in the controlled environment of liposome-based transport assays are needed to test this hypothesis.

hSLC19A3 interacts with many commonly used medical compounds. This has implications for the adjustment and monitoring of existing treatment plans, as well as the development of novel drugs^10,11^. In this work, we identified nine new high-affinity binders of the transporter, that are likely to inhibit thiamine uptake in the human body. Among the most potent ones are the tyrosine kinase inhibitors imatinib, gefitinib and dasatinib (Fig. 3). While these *in vitro* findings still need to be validated *in vivo*, our study highlights the importance of screening drug interactions of vitamin transporters like hSLC19A3. One aspect we did not address in the present study is the mode of inhibition of hSLC19A3 by the different drugs. For fedratinib and trimethoprim, it is known that they are xenobiotic substrates of the transporter^7^, and consequently inhibit thiamine uptake by acting as competing substrates. In this context it seems plausible that hSLC19A3 might also represent a transport route for existing and prospective medical compounds. In particular, it might be an interesting entry gate for drugs to cross the blood-brain barrier^7,49^. Our structural data provide a basis for the rational design of drugs to prevent the side effects of thiamine uptake inhibition or to use hSLC19A3 for tissue-specific drug delivery^49^. The other high-affinity thiamine transporter, hSLC19A2, shares high sequence identity (48%)^26^ and, based on AF2 predictions^39^, high structural conservation with hSLC19A3 (Supplementary Fig. 16b). We speculate that many of our findings also apply to hSLC19A2. It seems likely that this transporter follows a similar transport mechanism. Since the substrate binding site is well conserved, most inhibitors of hSLC19A3 are expected to bind to its closest relative.

With this study, we present a structural framework of thiamine transport and drug interactions of hSLC19A3. We provide structural snapshots of its transport cycle and detailed molecular insights into the binding of its substrate and high-affinity inhibitors to the transporter. This work can serve as a basis for studies of the transport mechanism of human MFS transporters and can guide future structure-based drug design approaches.

## Methods

### Cloning, expression and purification of hSLC19A3

The full-length, wildtype hSLC19A3 gene (Uniprot accession number: Q9BZV2) was cloned from cDNA into a modified pXLG vector^50^. In this vector, SLC19A3 is fused with a C-terminal human rhinovirus 3C (HRV-3C) protease cleavage site (LEVLFQGP), followed by a short flexible linker (SSG) and a Twin-Streptavidin tag for affinity purification. All mutants were generated using SLiCE cloning^51^. The transporter constructs were transiently transfected in Expi293F^TM^ cells using PEI MAX^®^ as a transfection agent (ratio 1:2 (w/w) of DNA to PEI MAX^®^) and grown for 96 hours post-transfection in FreeStyle^TM^ medium, at 37 °C, 8% (v/v) CO_2_, and 270 rpm^52^. Cells were harvested at 3,000×g, washed with 1× PBS (pH7.4) and subsequently frozen at −70°C. After thawing at room temperature, five grams of cell pellet were resuspended in 25 mL of solubilisation buffer (1×PBS, pH7.4, 200 mM NaCl, 5% glycerol (v/v), 1% LMNG (w/v), 0.1% CHS (w/v), 1 × EDTA-free protease inhibitors (Roche), 0.5 mM TCEP, 20 U/mL DNase I, 2.5 U avidin) in a 50 mL tube and incubated on a shaking platform for 1h at 4 °C. Cellular debris was removed via centrifugation for 30 min at 35,000×g at 4 °C. The supernatant was loaded onto 1 mL StrepTactin resin and incubated for 1 h at 4 °C. Afterwards, the resin was washed with 4 × 20 column volumes of StrepA buffer (20 mM Tris-HCl, pH 7.4, 350 mM NaCl, 5% glycerol (v/v), 0.5 mM TCEP, 0.02% LMNG (w/v), 0.002% CHS (w/v)). The target proteins were eluted in 5 column volumes StrepB buffer (20 mM Tris-HCl, pH 7.4, 150 mM NaCl, 0.002% LMNG, 0.0002% CHS, 10 mM desthiobiotin). The eluted protein was concentrated to 0.5 mL and further purified by size-exclusion chromatography (SEC) in SEC buffer (20 mM Tris-HCl, pH7.4, 150 mM NaCl, 0.002% LMNG (w/v), 0.0002% CHS (w/v)) on a Superdex S200 10/300 Increase column (Cytiva). Biotinylation of Avi-tagged protein for nanobody selection was performed by adding 120 μg/mL of BirA, 5 mM ATP, 50 μM biotin, 0.5 mM TCEP and 40 μg/mL 3C protease to the concentrated StrepB elution. The mixture was then incubated at 4 °C overnight. Subsequently, the transporter was separated from the enzymes and the small molecule additives by SEC as described above.

### Thermal stability measurements and binding affinity determination

The thermal stability of the hSLC19A3 constructs and their binding affinities for small molecules were determined by nanoDSF, using a Prometheus NT.48 (NanoTemper Technologies). For sample preparation, the protein was diluted to a final concentration of 4 μM in SEC buffer and supplied with the respective compounds in a concentration range from 2 μM - 2 mM. Before measurement, the samples were incubated for 30 min at room temperature. Melting scans were performed at a temperature range from 20 to 95 °C with a temperature increment of 1 °C per minute. Thermal unfolding curves were recorded by measuring the ratio of the tryptophan-fluorescence at 350 nm (F350) and 330 nm (F330). The protein melting temperature (T_m_) of a given sample was determined as the inflection point of the thermal unfolding curves, or the local maximum of its first derivative. The measured curves were analysed using the manufacturer’s software (PR.ThermControl, v 2.1.2, NanoTemper Technologies). Based on the determined T_m_ values, ligand induced thermal shifts (ΔT_m_) were determined as the difference between the melting temperature of the ligand-bound state (T_m_) and the apo state (T_m_^apo^). For the determination of apparent binding affinities at 25 °C, the T_m_ values recorded from dilution series of the respective ligands were used to fit the following equation:

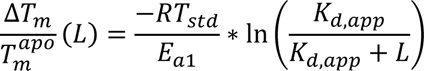

were L is the ligand concentration, R is the universal gas constant, T_std_ is the chosen standard temperature of 298.15 K (25 °C), E_a1_ is the activation energy for the unfolding of the apo state.

As R, T_std_ and L are known and the ΔT_m_ and T_m_^apo^ are experimentally determined, non-linear curve fitting can be used to estimate E_a1_ and K_d_,_app_. The fitting was performed using symfit (v. 0.5.3), as described in Kotov et al., 2023^53^, based on theoretical considerations formulated in Hall, 2019^54^.

### Generation of nanobodies

To generate nanobodies against the human thiamine transporter SLC19A3, the glycosylation-free mutant hSLC19A3-gf was injected into a llama (*Glama lama*) on a weekly basis over the course of six weeks as described elsewhere^28^. Following mRNA extraction and cDNA library preparation from peripheral blood mononuclear cells of the immunized llamas, VHH-fragments were cloned into phage display vectors. Subsequently, two rounds of phage display were performed to enrich for specific binders. From the enriched libraries, nanobody (VHH) families were identified by sequencing of 192 colonies. Eventually, three unique nanobodies, Nb3.3, Nb3.4 and Nb3.7 were identified and expressed. Binding to the transporters was confirmed using biolayer interferometry (Supplementary Fig. 2). For large scale production, the nanobodies were expressed in *E. coli* WK6 cells, using a PelB N-terminal signal peptide to direct them into the periplasm. After periplasmic extraction, the nanobodies were purified using nickel affinity chromatography, as previously described^28^.

### Biolayer interferometry (BLI)

Binding between hSLC19A3 and the nanobodies Nb3.3, Nb3.4 and Nb3.7 was measured by biolayer interferometry (BLI) using an Octet RED96 system (FortéBio). The nanobodies were immobilised at 300 nM via their C-terminal 6×His-tag on Octet® HIS1K Biosensors, pre-equilibrated in BLI buffer (20 mM Tris-HCl, pH7.4, 150 mM NaCl, 0.02% LMNG (w/v), 0.002% CHS (w/v), 0.1% BSA (w/v), 0.5 mM thiamine). After a baseline step of 60 s, the biosensors were dipped into BLI buffer containing hSLC19A3-gf at concentrations ranging from 25-800 nM for 240 s to allow for the binding of the transporter to the immobilised nanobodies. Afterwards, the sensors were transferred into fresh BLI buffer and the dissociation was followed over the course of 600 s. BLI experiments were performed at 22 °C. Data were reference-subtracted and aligned in the Octet Data Analysis software v10.0 (FortéBio). Binding affinities were determined using a 1:1 binding model and the maximum response values as readout.

### Cryo-EM sample preparation and data acquisition

For cryo-EM, hSLC15A3 purified in LMNG/CHS was used. For the inward-open apo state, as well as for the apo and thiamine-bound outward-open structures, wildtype hSLC19A3 was prepared in complex with Nb3.3 and Nb3.4, respectively. For all other ligand bound structures, the glycosylation-free hSLC19A3-gf in complex with Nb3.7 was used. After concentrating the transporter to 60 μM, the sample was supplied with a 1.5 × molar excess of nanobody and diluted to a total concentration of 30 μM of hSLC15A3 in SEC buffer. Compounds were added during the dilution step at the following concentrations: 500 μM thiamine, 200 μM hydroxychloroquine, 250 μM fedratinib, 300 μM amprolium or 250 μM amitriptyline. The samples were subsequently incubated for 4-18 hours on ice. 3.6 μL of sample was applied to glow discharged holey carbon film grids (either Quantifoil 300 mesh, Au, R1.2/1.3, or R2/1, or C-flat, 200 mesh, Cu, 2/1) for 15 seconds at 10 °C and 100% humidity in a Vitrobot Mark IV (Thermo Fisher Scientific). Blotting was performed for 3.5 s at blot force −5 and with a drain time of 0.5 s. The grids were plunged into liquid ethane/propane and transferred to liquid nitrogen for storage. Data were collected in counting mode on a Titan Krios G3 (FEI) operating at 300 kV with a BioQuantum imaging filter (Gatan) and a K3 direct detection camera (Gatan). Data was collected in fast acquisition mode by aberration-free image shift (AFIS) at 105,000 × magnification, a physical pixel size of 0.83 or 0.85 Å, a slit width of 20 eV and a total dose of 45-60 e^−^/Å^2^ (dose rate of ∼15 e^−^/px/s, exposure time of ∼2.4 s; for details, see table 1). The nominal defocus was for most datasets was set to the range of −0.8 μM to −1.6 μm.

### Cryo-EM data processing

Processing of the acquired micrographs was performed using cryoSPARC (v. 4.4)^55^. Briefly, the collected movies were motion corrected using patch motion correction. After patch CTF estimation, exposures with a CTF resolution <3.5 Å, a relative ice thickness <1.05 and a total frame motion distance <30 pixels were selected for further processing. Particle picking was performed using template picker. The corresponding templates were generated from blob-picking (size of blob) a small subset of the data (200 - 600 micrographs), which was subjected to 2D classification, *ab-initio* reconstruction and non-uniform refinement (NU-refinement). After template-based particle picking of the entire dataset, protein-containing and well-aligning particles were enriched using several rounds of 2D classification. The selected particles were then used in a sequence of *ab-initio* reconstruction and NU-refinement to create a first higher resolution map, usually in the resolution range of 3.5-4.0 Å. This refined map was then, together with randomly generated volumes, used for several rounds of heterogeneous refinement to retrieve more good particles and remove junk particles. The resulting map was then further refined using NU-refinement. The corresponding particles were subsequently subjected to local motion correction and re-extracted with a larger box size of 384 pixels (∼326 Å). After another round of NU-refinement, the resulting density map was locally refined using a mask covering the transporter-nanobody complex. This was followed by local resolution estimation and local filtering to finalise the maps. For more details, see Supplementary Fig. 5.

### Model building and refinement

For starting models, AlphaFold2 predictions of hSLC19A3 and the nanobodies were used^39^. These were relaxed into the corresponding EM density maps using ISOLDE^56^. Ligands were built and parametrised using the eLBOW package within PHENIX^57^ and integrated in the protein structures using Coot (v. 0.9.8.1)^58^. The resulting models were subsequently refined iteratively in PHENIX and Coot. Figures were prepared using UCSF ChimeraX (v. 1.6.1)^56^.

## Data Availability

The EM data and fitted models for hSLC19A3 have been deposited in the Electron Microscopy Data Bank and the PDB under the following accession codes: hSLC19A3-wt:Nb3.4-apo (8S4U.pdb, EMD-19716); hSLC19A3-wt:Nb3.4:thiamine (8S5U.pdb, EMD-19750); hSLC19A3-wt:Nb3.3-apo (8S5V.pdb, EMD-19751); hSLC19A3-gf:Nb3.7:thiamine (8S61.pdb, EMD-19754); hSLC19A3-gf:Nb3.7:Fedratinib (8S5W.pdb, EMD-19752); hSLC19A3-gf:Nb3.7:Amprolium (8S62.pdb, EMD-19755), hSLC19A3-gf:Nb3.7: Hydroxy-chloroquine (8S5Z.pdb, EMD-19753); hSLC19A3-gf:Nb3.7:Amitriptyline (8S66.pdb, EMD-19756)

## Materials Availability

All reagents generated in this study are available from the Lead Contact with a completed Materials Transfer Agreement.

## Acknowledgements

We thank the Sample Preparation and Characterization facility at EMBL (Hamburg, Germany) and the team of the cryo-EM Facility at CSSB for their support, technical assistance and advice. We want to acknowledge Tânia Custódio and all group members for fruitful discussions and continuous support and feedback on the project. Part of this work was performed at the CryoEM Facility at CSSB, supported by the UHH and DFG (grant numbers INST 152/772-1|152/774-1|152/775-1|152/776-1|152/777-1 FUGG). Nanobody discovery was realised through the Nanobodies4Instruct centre and the support by Instruct-ERIC (PID: 23688), which is part of the European Strategy Forum on Research Infrastructures (ESFRI). We want to thank in particular Alison Lundqvist and Eva Beke from the Steyaert Lab at VUB for their technical assistance during nanobody discovery. The research-stay of F.G. at VUB was supported by an EMBO Scientific Exchange Grand (Nr. 10251).

## Author Contributions

F.G. conceptualised the project and experiments, cloned and expressed transporter constructs and nanobodies, performed the nanobody discovery, performed and analysed nanoDSF and BLI experiments, collected and processed all cryo-EM data and prepared the manuscript. L.S. and A.F. cloned and expressed transporter constructs, as well as nanobodies, and collected biophysical data. K.E.J.J. and S.M. helped with experiment design, as well as cryo-EM data processing, structure refinement and manuscript writing. J.S and E.P. immunised llamas and generated nanobody libraries for the discovery process. C.L. was responsible for the overall project administration, experiment design, funding acquisition, and for reviewing and editing of the manuscript.

## References

1. Bettendorff L. Thiamin. In: Erdman Jr JW, McDonald IA, Zeisel SH, E. Present Knowledge in Nutrition - Thiamin. in 261–79. (2012).

2. Timm, D. E., Liu, J., Baker, L. J. & Harris, R. A. Crystal structure of thiamin pyrophosphokinase. J. Mol. Biol. 310, 195–204 (2001).

3. Baker, L. J., Dorocke, J. A., Harris, R. A. & Timm, D. E. The crystal structure of yeast thiamin pyrophosphokinase. Structure 9, 539–546 (2001).

4. Tylicki, A., Lotowski, Z., Siemieniuk, M. & Ratkiewicz, A. Thiamine and selected thiamine antivitamins — biological activity and methods of synthesis. Biosci. Rep. 38, 1–23 (2018).

5. Zhao, R. & Goldman, I. D. Folate and thiamine transporters mediated by facilitative carriers (SLC19A1-3 and SLC46A1) and folate receptors. Mol. Aspects Med. 34, 373–385 (2013).

6. Dutta, B. et al. Cloning of the human thiamine transporter, a member of the folate transporter family. J. Biol. Chem. 274, 31925–31929 (1999).

7. Giacomini, M. M. et al. Interaction of 2,4-diaminopyrimidine-containing drugs including fedratinib and trimethoprim with thiamine transporters. Drug Metab. Dispos. 45, 76–85 (2017).

8. Yamashiro, T., Yasujima, T., Said, H. M. & Yuasa, H. PH-dependent pyridoxine transport by SLC19A2 and SLC19A3: Implications for absorption in acidic microclimates. J. Biol. Chem. 295, 16998–17008 (2020).

9. Chen, L. et al. OCT1 is a high-capacity thiamine transporter that regulates hepatic steatosis and is a target of metformin. Proc. Natl. Acad. Sci. U. S. A. 111, 9983–9988 (2014).

10. Vora, B. et al. Drug-nutrient interactions: Discovering prescription drug inhibitors of the thiamine transporter ThTR-2 (SLC19A3). Am. J. Clin. Nutr. 111, 110–121 (2020).

11. Wen, A. et al. The Impacts of Slc19a3 Deletion and Intestinal SLC19A3 Insertion on Thiamine Distribution and Brain Metabolism in the Mouse. Metabolites 13, (2023).

12. Reidling, J. C., Lambrecht, N., Kassir, M. & Said, H. M. Impaired Intestinal Vitamin B1 (Thiamin) Uptake in Thiamin Transporter-2-Deficient Mice. Gastroenterology 138, 1802–1809 (2010).

13. Kato, K. et al. Involvement of organic cation transporters in the clearance and milk secretion of thiamine in mice. Pharm. Res. 32, 2192–2204 (2015).

14. Kono, S. et al. Mutations in a Thiamine-Transporter Gene and Wernicke’s-like Encephalopathy. N. Engl. J. Med. 360, 1792–1794 (2009).

15. Wang, J. et al. Report of the Largest Chinese Cohort With SLC19A3 Gene Defect and Literature Review. Front. Genet. 12, 1–9 (2021).

16. Zeng, W. Q. et al. Biotin-responsive basal ganglia disease maps to 2q36.3 and is due to mutations in SLC19A3. Am. J. Hum. Genet. 77, 16–26 (2005).

17. Alfadhel, M. et al. Targeted SLC19A3 gene sequencing of 3000 Saudi newborn: a pilot study toward newborn screening. Ann. Clin. Transl. Neurol. 6, 2097–2103 (2019).

18. Pardanani, A. et al. Safety and efficacy of fedratinib in patients with primary or secondary myelofibrosis: A randomized clinical trial. JAMA Oncol. 1, 643–651 (2015).

19. Zhang, Q. et al. The Janus kinase 2 inhibitor fedratinib inhibits thiamine uptake: A putative mechanism for the onset of Wernicke’s encephalopathy. Drug Metab. Dispos. 42, 1656–1662 (2014).

20. Quistgaard, E. M., Löw, C., Guettou, F. & Nordlund, P. Understanding transport by the major facilitator superfamily (MFS): Structures pave the way. Nat. Rev. Mol. Cell Biol. 17, 123–132 (2016).

21. Drew, D., North, R. A., Nagarathinam, K. & Tanabe, M. Structures and General Transport Mechanisms by the Major Facilitator Superfamily (MFS). Chem. Rev. 121, 5289–5335 (2021).

22. Hou, Z. & Matherly, L. H. Biology of the major facilitative folate transporters SLC19A1 and SLC46A1. Curr. Top. Membr. 73, 175–204 (2014).

23. Said, H. M., Balamurugan, K., Subramanian, V. S. & Marchant, J. S. Expression and functional contribution of hTHTR-2 in thiamin absorption in human intestine. Am. J. Physiol. - Gastrointest. Liver Physiol. 286, 491–498 (2004).

24. Liang, X. et al. Metformin Is a Substrate and Inhibitor of the Human Thiamine Transporter, THTR-2 (SLC19A3). Mol. Pharm. 12, 4301–4310 (2015).

25. Subramanian, V. S., Marchant, J. S. & Said, H. M. Biotin-responsive basal ganglia disease-linked mutations inhibit thiamine transport via hTHTR2: Biotin is not a substrate for hTHTR2. Am. J. Physiol. - Cell Physiol. 291, 851–859 (2006).

26. Rajgopal, A., Edmondnson, A., Goldman, I. D. & Zhao, R. SLC19A3 encodes a second thiamine transporter ThTr2. Biochim. Biophys. Acta - Mol. Basis Dis. 1537, 175–178 (2001).

27. Miyake, K., Yasujima, T., Takahashi, S., Yamashiro, T. & Yuasa, H. Identification of the amino acid residues involved in the species-dependent differences in the pyridoxine transport function of SLC19A3. J. Biol. Chem. 298, 102161 (2022).

28. Pardon, E. et al. A general protocol for the generation of Nanobodies for structural biology. Nat. Protoc. 9, 674–693 (2014).

29. Zhang, Q. et al. Recognition of cyclic dinucleotides and folates by human SLC19A1. Nature 612, 170–176 (2022).

30. Wright, N. J. et al. Methotrexate recognition by the human reduced folate carrier SLC19A1. Nature 609, (2022).

31. Dang, Y. et al. Molecular mechanism of substrate recognition by folate transporter SLC19A1. Cell Discov. 8, (2022).

32. Drew, D. & Boudker, O. Shared Molecular Mechanisms of Membrane Transporters. Annu. Rev. Biochem. 85, 543–572 (2016).

33. Harris, T. K. & Mildvan, A. S. High-precision measurement of hydrogen bond lengths in proteins by nuclear magnetic resonance methods. Proteins Struct. Funct. Genet. 35, 275–282 (1999).

34. Bas, D. C., Rogers, D. M. & Jensen, J. H. Very fast prediction and rationalization of pKa values for protein-ligand complexes. Proteins Struct. Funct. Genet. 73, 765–783 (2008).

35. Zhao, R., Assaraf, Y. G. & Goldman, I. D. A mutated murine reduced folate carrier (RFC1) with increased affinity for folic acid, decreased affinity for methotrexate, and an obligatory anion requirement for transport function. J. Biol. Chem. 273, 19065–19071 (1998).

36. Yee, S. W. et al. SLC19A1 Pharmacogenomics Summary. Pharmacogenet Genomics 20, 708–715 (2010).

37. Labay, V. et al. Mutations in SLC19A2 cause thiamine-responsive megaloblastic anaemia associated with diabetes mellitus and deafness. Nat. Genet. 22, 300–304 (1999).

38. Liberman, M. C., Tartaglini, E., Fleming, J. C. & Neufeld, E. J. Deletion of SLC19A2, the high affinity thiamine transporter, causes selective inner hair cell loss and an auditory neuropathy phenotype. JARO - J. Assoc. Res. Otolaryngol. 7, 211–217 (2006).

39. Jumper, J. et al. Highly accurate protein structure prediction with AlphaFold. Nature 596, 583–589 (2021).

40. Zeng, Y. C. et al. Structural basis of promiscuous substrate transport by Organic Cation Transporter 1. (2023) doi:10.1038/s41467-023-42086-9.

41. Ozand, P. T. et al. Biotin-responsive basal ganglia disease: A novel entity. Brain 121, 1267–1279 (1998).

42. Aburezq, M. et al. Biotin-thiamine responsive basal ganglia disease: a retrospective review of the clinical, radiological and molecular findings of cases in Kuwait with novel variants. Orphanet J. Rare Dis. 18, 1–13 (2023).

43. Wesół-Kucharska, D., et al. Early treatment of biotin–thiamine–responsive basal ganglia disease improves the prognosis. Mol. Genet. Metab. Reports 29, (2021).

44. Kevelam, S. H. et al. Exome sequencing reveals mutated SLC19A3 in patients with an early-infantile, lethal encephalopathy. Brain 136, 1534–1543 (2013).

45. Rose-Sperling, D., Tran, M. A., Lauth, L. M., Goretzki, B. & Hellmich, U. A. 19 F NMR as a versatile tool to study (membrane) protein structure and dynamics. Biol. Chem. 400, 1277–1288 (2019).

46. Aljabri, M. F., Kamal, N. M., Arif, M., AlQaedi, A. M. & Santali, E. Y. M. A case report of biotin–thiamine-responsive basal ganglia disease in a Saudi child. Medicine (Baltimore). 95, e4819 (2016).

47. Schänzer, A. et al. Stress-induced upregulation of SLC19A3 is impaired in biotin-thiamine-responsive basal ganglia disease. Brain Pathol. 24, 270–279 (2014).

48. Dudeja, P. K., Tyagi, S., Kavilaveettil, R. J., Gill, R. & Said, H. M. Mechanism of thiamine uptake by human jejunal brush-border membrane vesicles. Am. J. Physiol. - Cell Physiol. 281, 786–792 (2001).

49. Galetin, A. et al. Membrane transporters in drug development and as determinants of precision medicine. Nat. Rev. Drug Discov. (2024) doi:10.1038/s41573-023-00877-1.

50. Backliwal, G. et al. Rational vector design and multi-pathway modulation of HEK 293E cells yield recombinant antibody titers exceeding 1 g/l by transient transfection under serum-free conditions. Nucleic Acids Res. 36, (2008).

51. Zhang, Y., Uwe Werling, W. & Edelmann, W. Seamless Ligation Cloning Extract (SLiCE) Cloning Method Yongwei. Methods Mol. Biol. 1116, 235–244 (2014).

52. Pieprzyk, J., Pazicky, S. & Löw, C. Transient expression of recombinant membrane-eGFP fusion proteins in HEK293 cells. Methods Mol. Biol. 1850, 17–31 (2018).

53. Kotov, V. et al. Plasticity of the binding pocket in peptide transporters underpins promiscuous substrate recognition. Cell Rep. 42, (2023).

54. Hall, J. A simple model for determining affinity from irreversible thermal shifts. Protein Sci. 28, 1880–1887 (2019).

55. Punjani, A., Rubinstein, J. L., Fleet, D. J. & Brubaker, M. A. CryoSPARC: Algorithms for rapid unsupervised cryo-EM structure determination. Nat. Methods 14, 290–296 (2017).

56. Pettersen, E. F. et al. UCSF ChimeraX: Structure visualization for researchers, educators, and developers. Protein Sci. 30, 70–82 (2021).

57. Afonine, P. V. et al. Real-space refinement in PHENIX for cryo-EM and crystallography. Acta Crystallogr. Sect. D Struct. Biol. 74, 531–544 (2018).

58. Emsley, P., Lohkamp, B., Scott, W. G. & Cowtan, K. Features and development of Coot. Acta Crystallogr. Sect. D Biol. Crystallogr. 66, 486–501 (2010).

