## Supplementary file for "Structural basis of substrate transport and drug recognition by the human thiamine transporter SLC19A3"

### Supplementary Information

### Supplementary Figures

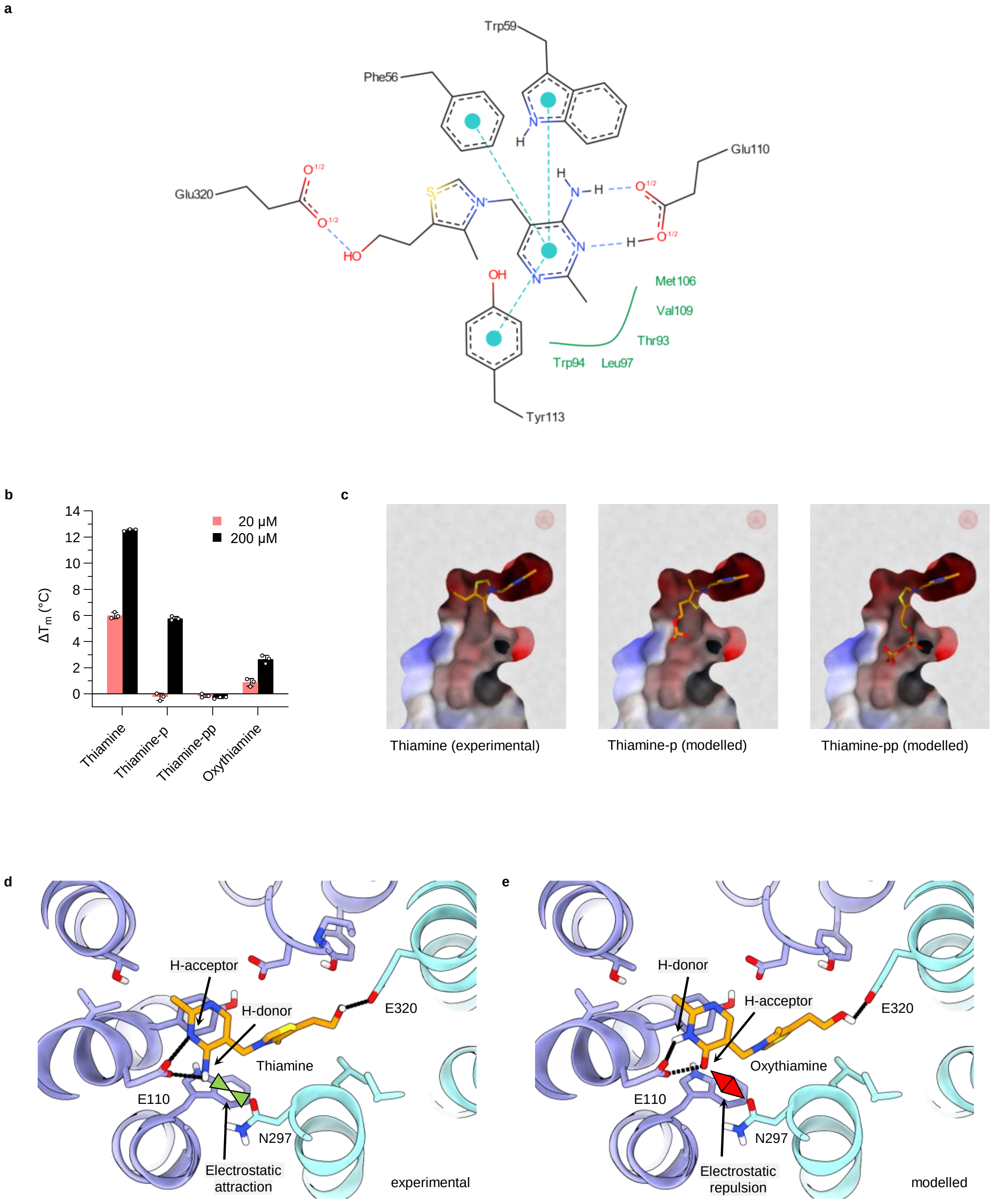


**Supplementary Fig. 1: Recognition of thiamine.**

**Supplementary Fig. 1: Recognition of thiamine. a**, Schematic representation of the coordination of thiamine by hSLC19A3. π-π interactions are indicated in cyan, hydrophobic contacts in green, hydrogen bonds in light blue. The interaction of the hydroxyethyl tail with Glu320 is only present in the inward-open state. In the outward-open state, this moiety forms polar contacts with the backbone of Ile36. **b**, Thermal shifts induced by the indicated compounds (fluorescence readout of thermal unfolding curve at 350 nm). While thiamine induces a strong stabilisation of hSLC19A3, its phosphorylated derivatives thiamine monophosphate (thiamine-p) shows a strongly reduced affinity for the transporter; and thiamine pyrophosphate (thiamine-pp) does not stabilise the transporter at all. The closely related derivative oxythiamine also stabilises the transporter only weakly. **c**, Structural model for the interaction of the phosphorylated derivatives of thiamine with hSLC19A3 in the inward-open state. The left panel shows the experimentally determined structure of hSLC19A3:Nb3.7:thiamine. The two panels to the right highlight thiamine-p and -pp modelled in the same EM density map using Coot. These models demonstrate the electrostatic mismatch between the negatively charged vestibule of the transporter and the negatively charged phosphate moieties. Furthermore, the important hydrogen bond between the hydroxyethyl tail of thiamine and Glu320 is disrupted by the phosphorylation of thiamine. **d**, Coordination of thiamine in hSLC19A3 in the inward-open state (hSLC19A3:Nb3.7:thiamine). The green arrows indicate the electrostatic attraction between the positively polarised hydrogens of the amino group of thiamine and the side chain carbonyl oxygen of Asn297. **e**, Oxythiamine modelled in the same EM density map. While the hydrogen bonding pattern with the transporter is in principle satisfied, the type of hydrogen bond pairs is different and might be energetically less favourable. Additionally, the carbonyl oxygen of oxythiamine is in electrostatic conflict with the side chain of Asn297. This might explain the observed weak interaction of this compound with hSLC19A3.

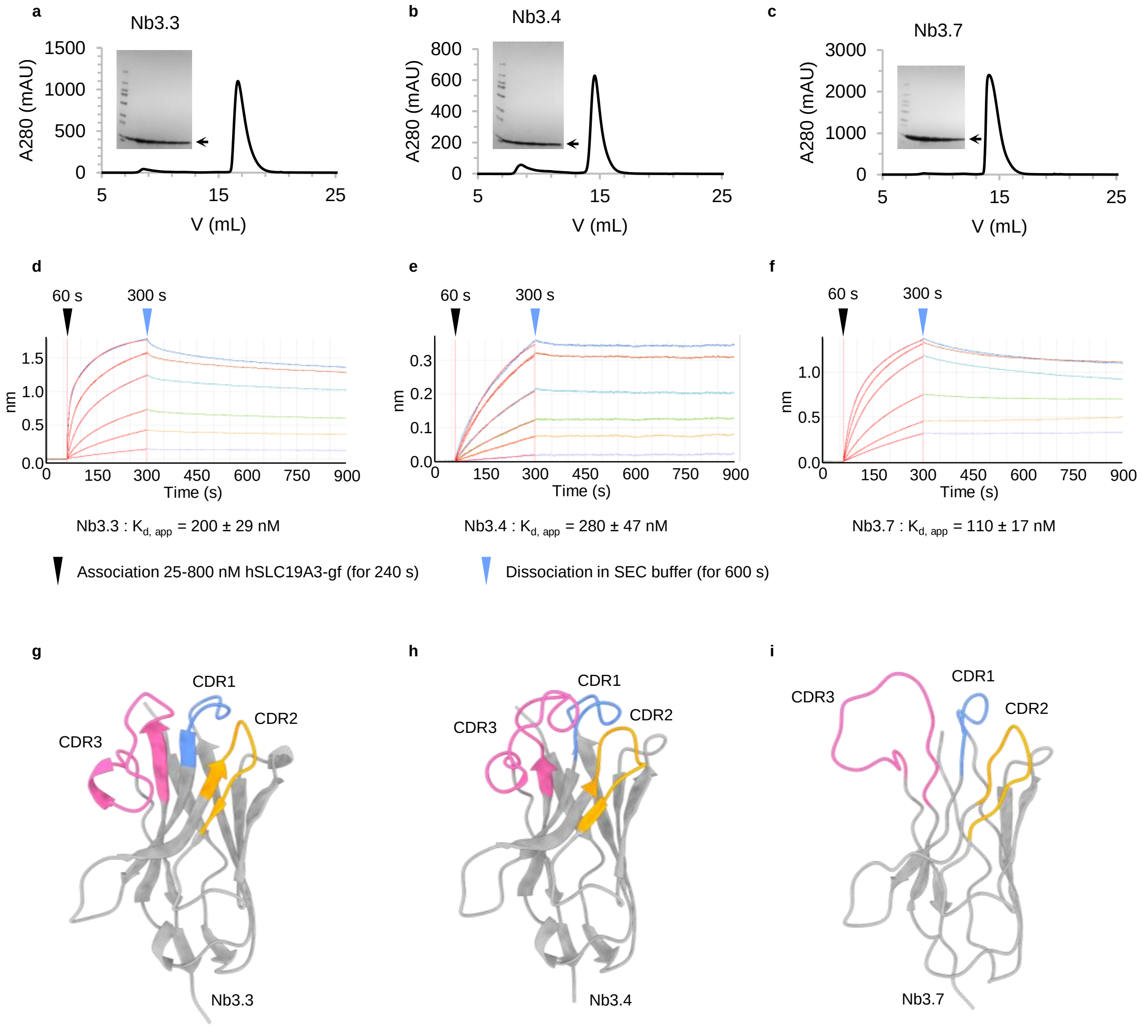


**Supplementary Fig. 2: Conformation-specific nanobodies against hSLC19A3. a-c**, Purification of the nanobodies Nb3.3, Nb3.4, and Nb3.7. Shown are size-exclusion chromatography (SEC) traces and SDS-PAGE gels of different fractions of the SEC peaks. **d-f**, Biolayer interferometry (Octet®) measurements of binding of hSLC19A3-gf to the respective nanobody immobilised on anti-penta-His sensors (Octet® HIS1 Biosensors). The sensors were incubated with different concentrations of hSLC19A3-gf; 25 nM (purple), 50 nM (yellow), 100 nM (green), 200 nM (light blue), 400 nM (orange), 800 nM (dark blue). Apparent binding affinities were determined by fitting the maximal response to a one-site binding model. **g-i**, Cryo-EM structures of the Nb3.3 and Nb3.4 when bound to hSLC19A3-wt and Nb3.7 bound to hSLC19A3-gf. The paratopes for the interaction with the transporter are formed by the highly variable CDRs (colour coded).

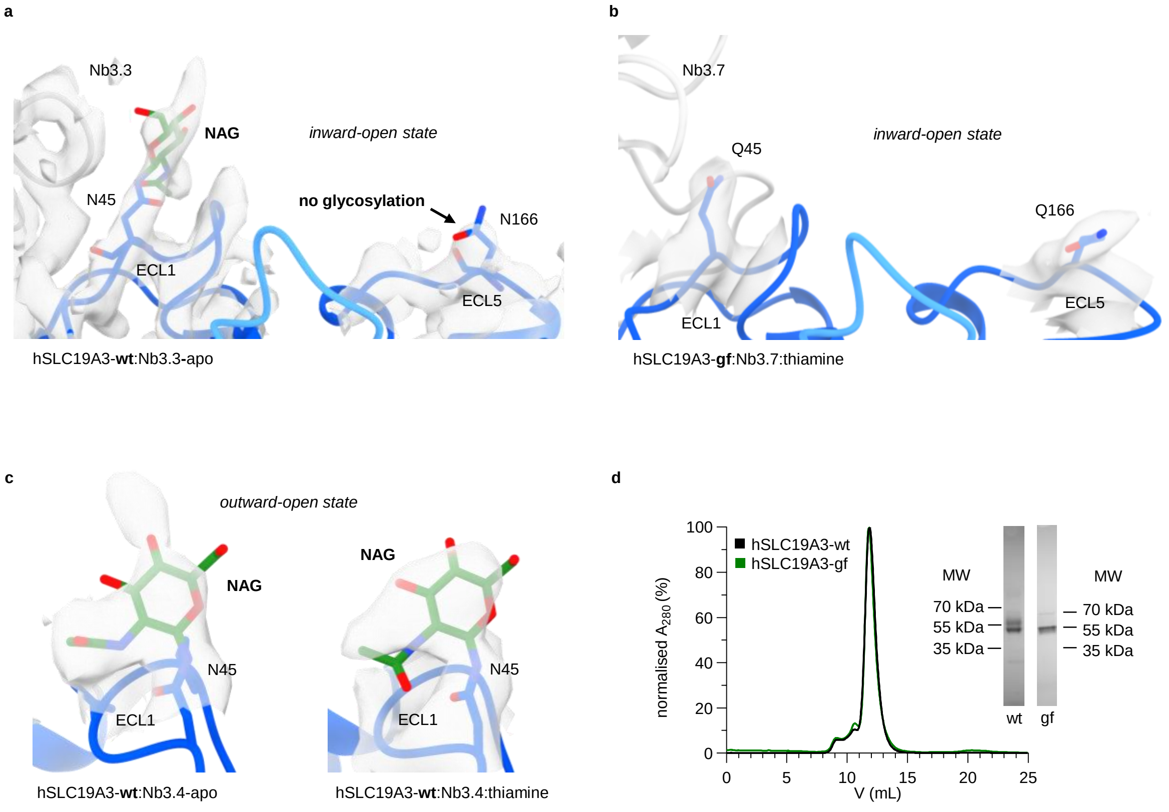


**Supplementary Fig. 3: N-linked glycosylation of hSLC19A3.** **a**, Structure model and cryo-EM density of hSLC19A3-wt:Nb3.3-apo. The wildtype transporter shows additional non-protein density extending from Asn45, but not from Asn166 (the two predicted N-glycosylation sites; map contour level = 1.01) indicative for a glycosylation site. **b**, For the hSLC19A3-gf double mutant (Asn45Gln, Asn166Gln), no extra density is visible. This confirms that the mutation of Asn45 to glutamine prevents N-linked glycosylation of the transporter (map contour level = 0.126). **c**, Glycan densities can be observed on Asn45 in the apo (left) and thiamine-bound (right) outward-open state of wildtype hSLC19A3 (map contour levels = 0.432 and 0.625, respectively). In these structures, there is likewise no extra density on Asn166, indicating that this residue is actually not glycosylated in recombinantly expressed hSLC19A3. **d**, Size-exclusion chromatography (SEC) trace of wildtype hSLC19A3 (wt) and the glycosylation-free hSLC19A3-gf double mutant (gf). The SEC traces are almost identical. SDS-PAGE (insets) show that the higher molecular weight band (slightly above 55 kDa) corresponding to the glycosylated transporter, is absent for hSLC19A3-gf.

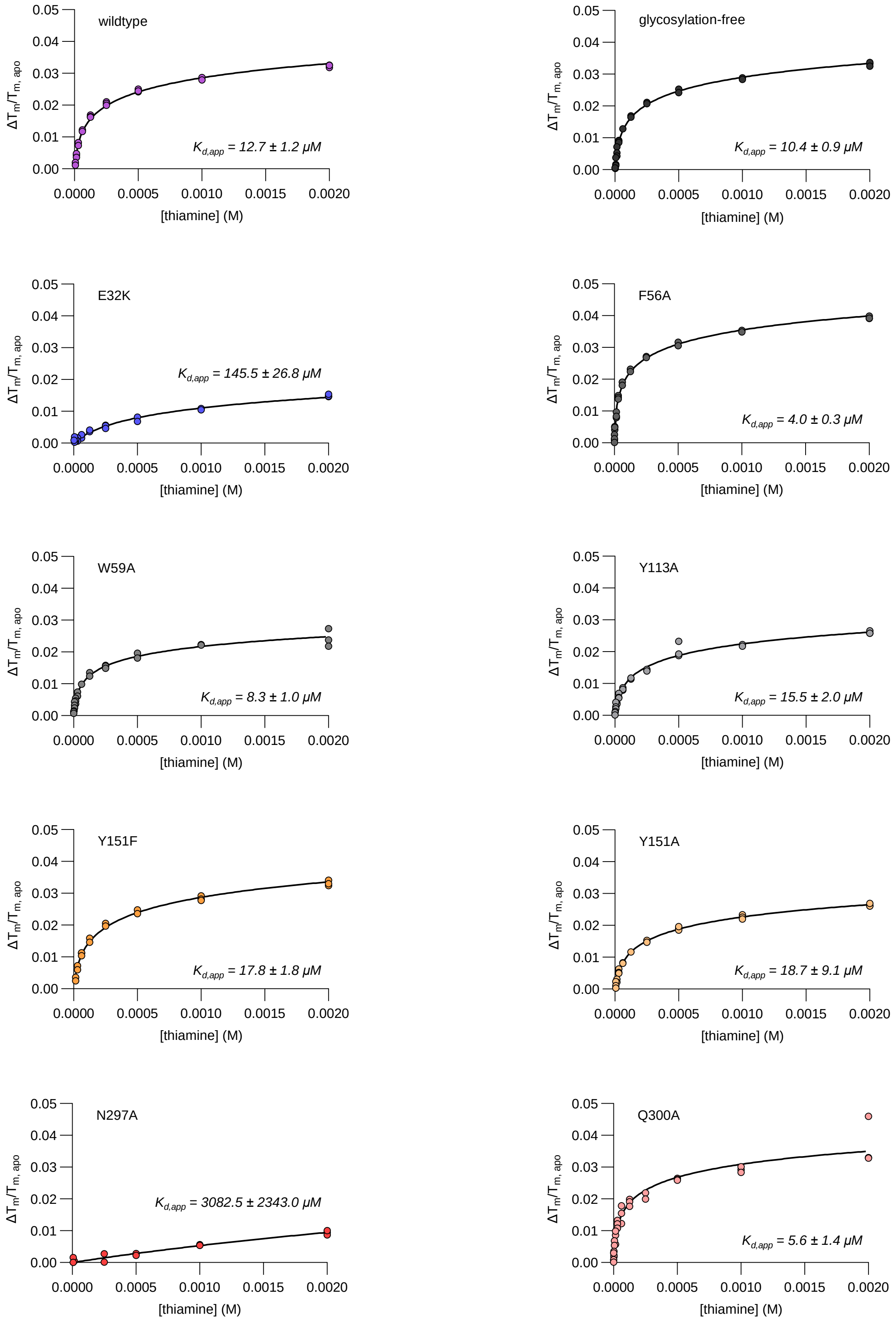


**Supplementary Fig. 4: Thiamine binding affinity of single point mutants of hSLC19A3.** The graphs highlight the measured thermostability shifts of the respective hSLC19A3 mutants in dependence of thiamine concentration (coloured circles, *n = 3*, independent measurements). The corresponding melting curves were read out based on the ratio of the fluorescence at 350 nm and 330 nm (F350/F330). The depicted curves were fitted as described in the methods section to determine the apparent dissociation constants $K_{d,app}$.

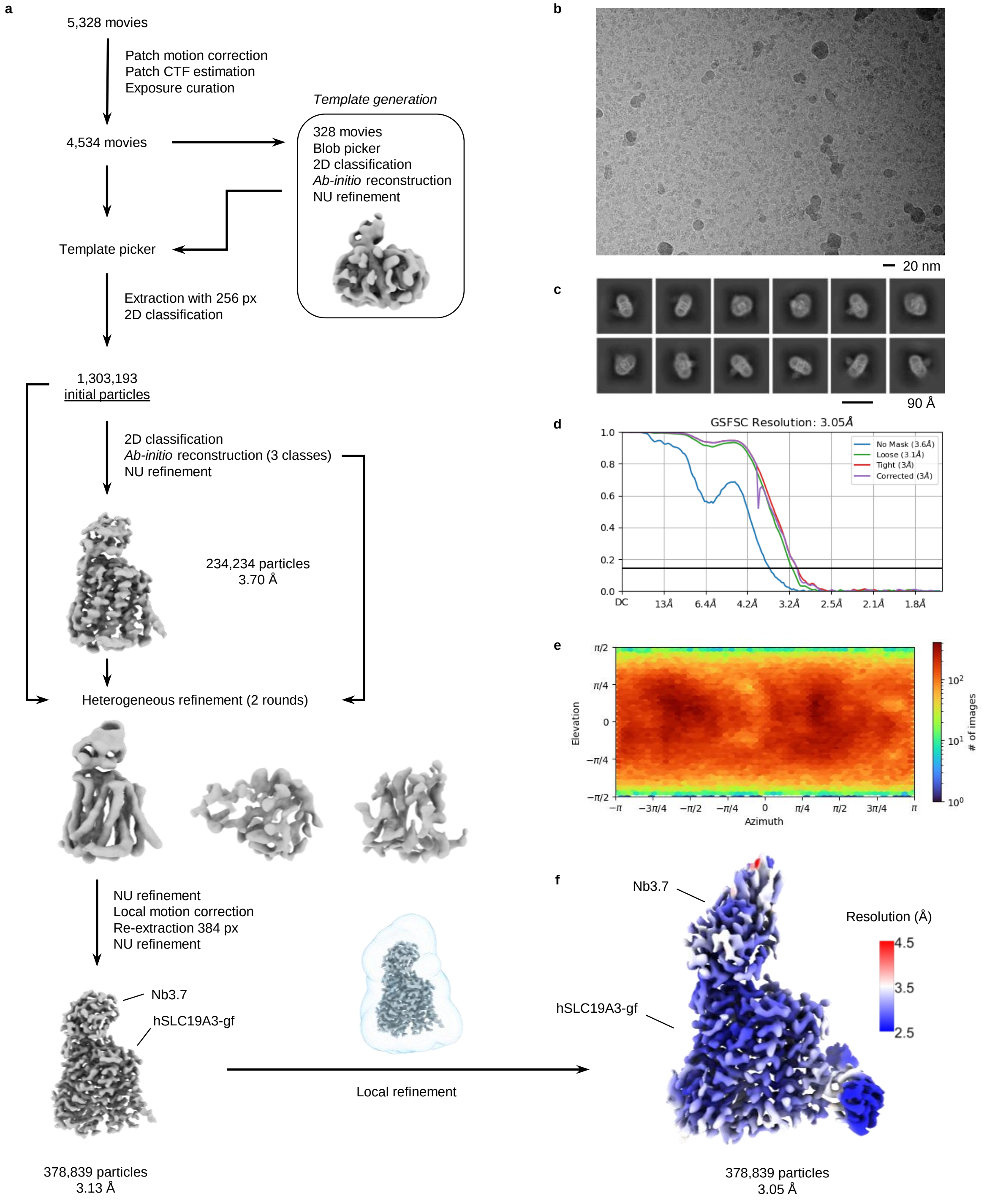


**Supplementary Fig. 5: Representative cryo-EM processing workflow.** **a**, Image processing workflow of the hSLC19A3:Nb3.7:fedratinib data set. The shown workflow was used with minor deviations for the processing of all presented cryo-EM data sets. **b**, Representative micrograph acquired at a defocus of -1.6 µm. **c**, 2D class averages. **d**, Gold-standard Fourier shell correlation (GSFSC) curves between two half maps. **e**, Angular distribution of particles in the final 3D local refinement. **f**, Cryo-EM density map of hSLC19A3:Nb3.7:fedratinib, surface coloured by local resolution.

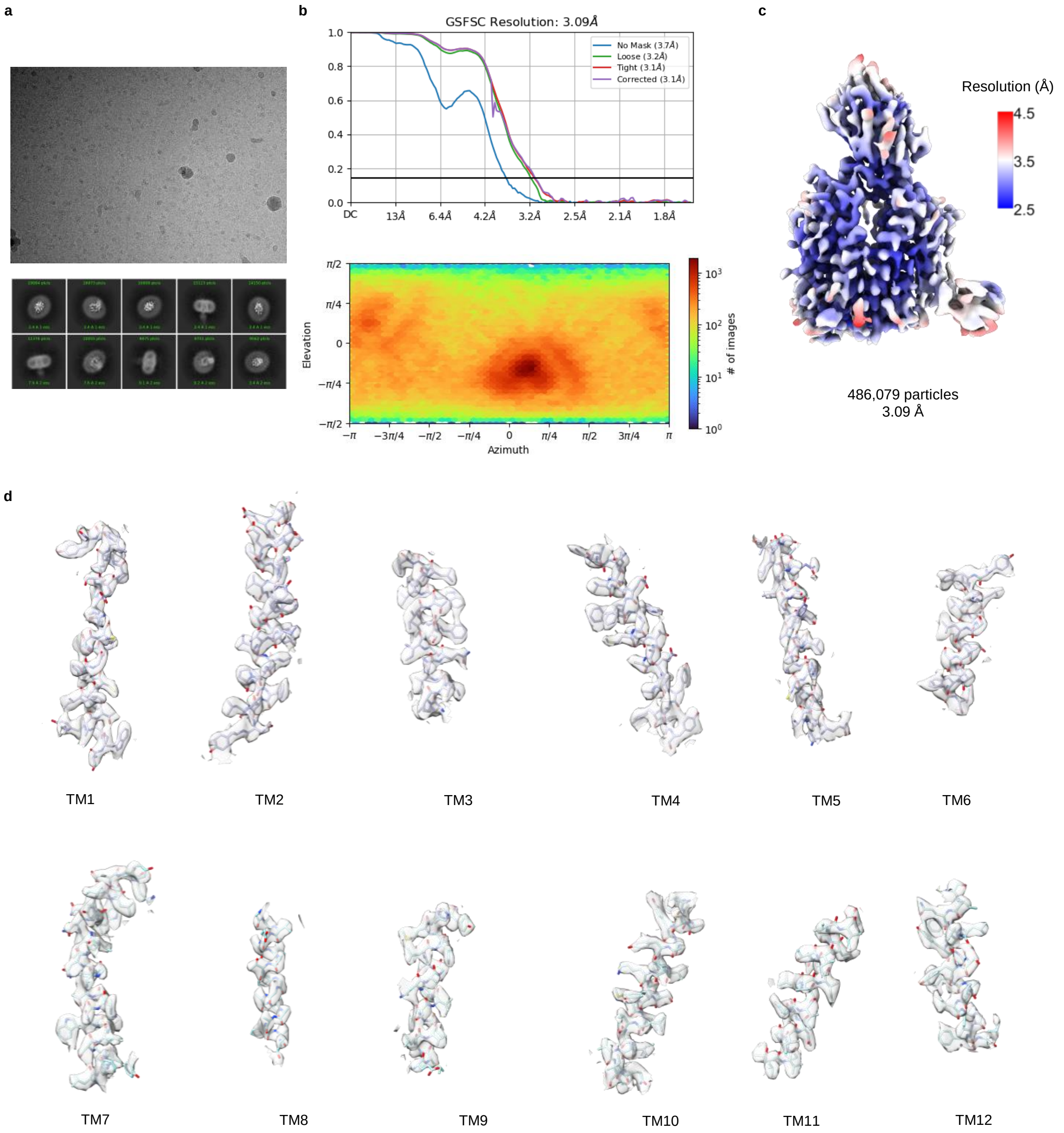


**Supplementary Fig. 6: Cryo-EM data processing overview for hSLC19A3-wt:Nb3.4-apo. a**, Representative micrograph (top) and 2D class averages (below). **b**, Gold-standard Fourier shell correlation (GSFSC) curves between two half maps (top). Angular distribution of particles in the final 3D local refinement (below). **c**, Local resolution estimation of the final cryo-EM map (map contour level = 0.57). **d**, Close-up on the density for the individual transmembrane helices (map contour level = 0.711).

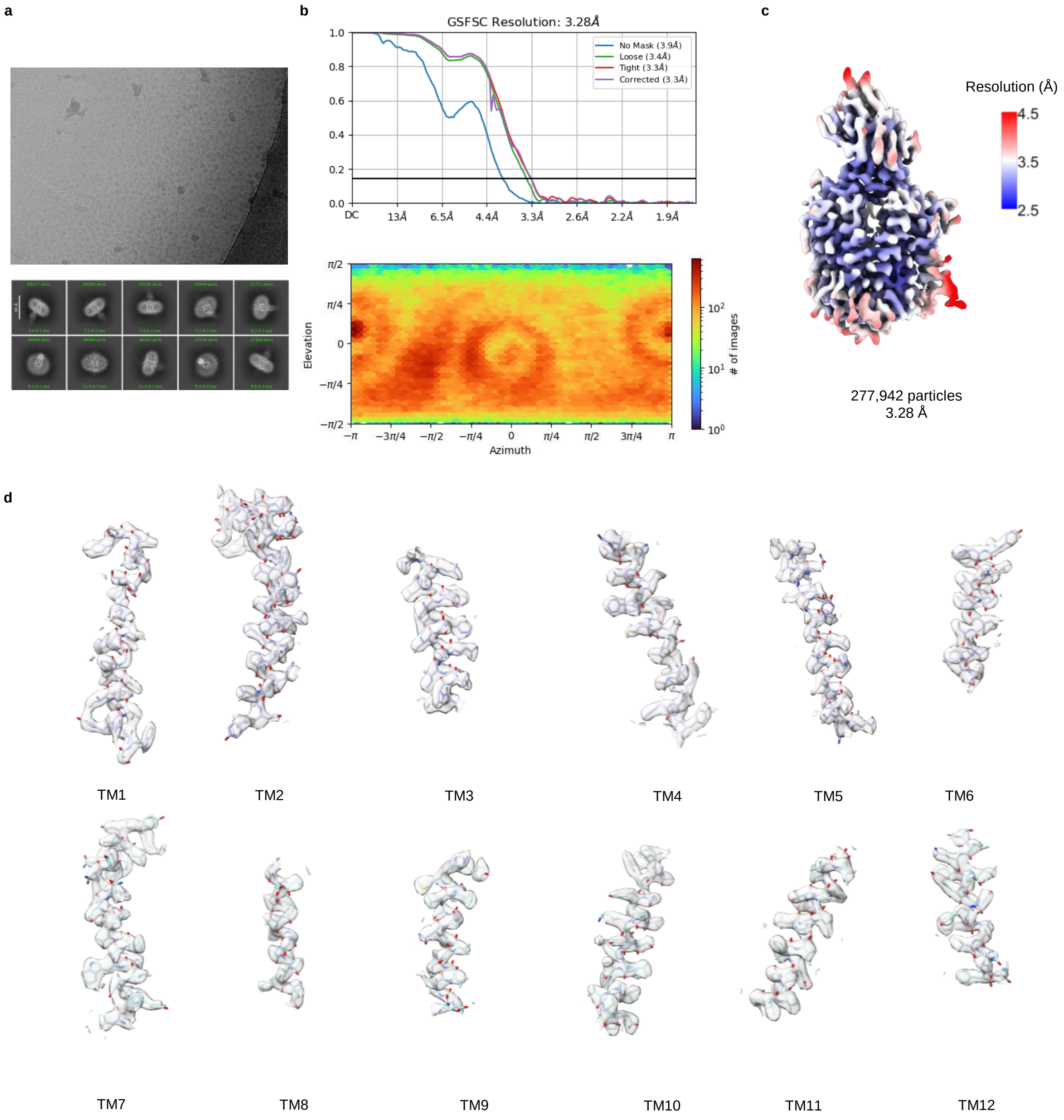


**Supplementary Fig. 7: Cryo-EM data processing overview for hSLC19A3-wt:Nb3.4:thiamine. a**, Representative micrograph (top) and 2D class averages (below). **b**, Gold-standard Fourier shell correlation (GSFSC) curves between two half maps (top). Angular distribution of particles in the final 3D local refinement (below). **c**, Local resolution estimation of the final cryo-EM map (map contour level = 0.676). **d**, Close-up on the density for the individual transmembrane helices (map contour level = 1.01).

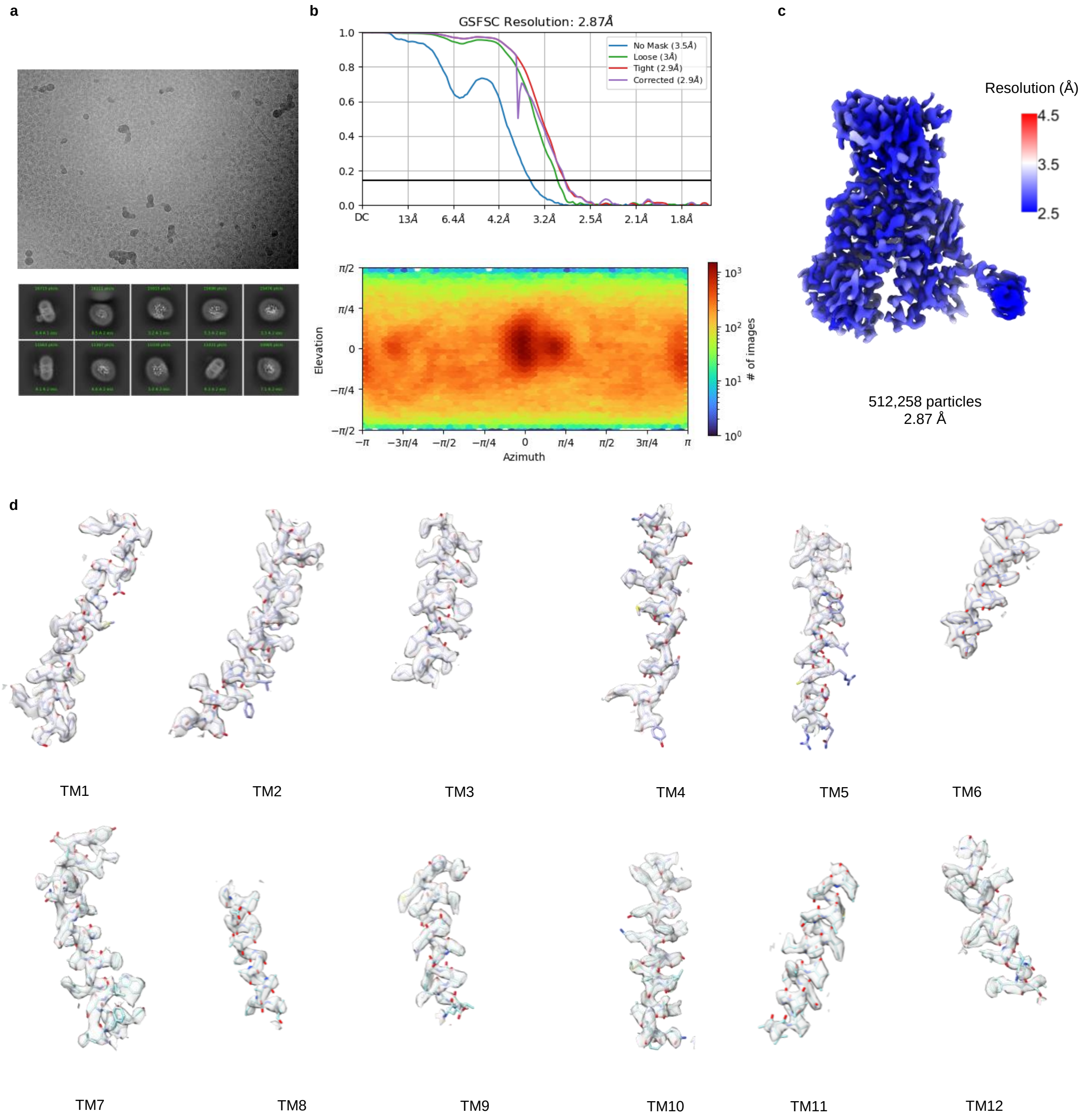


**Supplementary Fig. 8: Cryo-EM data processing overview for hSLC19A3-wt:Nb3.3-apo. a**, Representative micrograph (top) and 2D class averages (below). **b**, Gold-standard Fourier shell correlation (GSFSC) curves between two half maps (top). Angular distribution of particles in the final 3D local refinement (below). **c**, Local resolution estimation of the final cryo-EM map (map contour level = 0.951). **d**, Close-up on the density for the individual transmembrane helices (map contour level = 1.07).

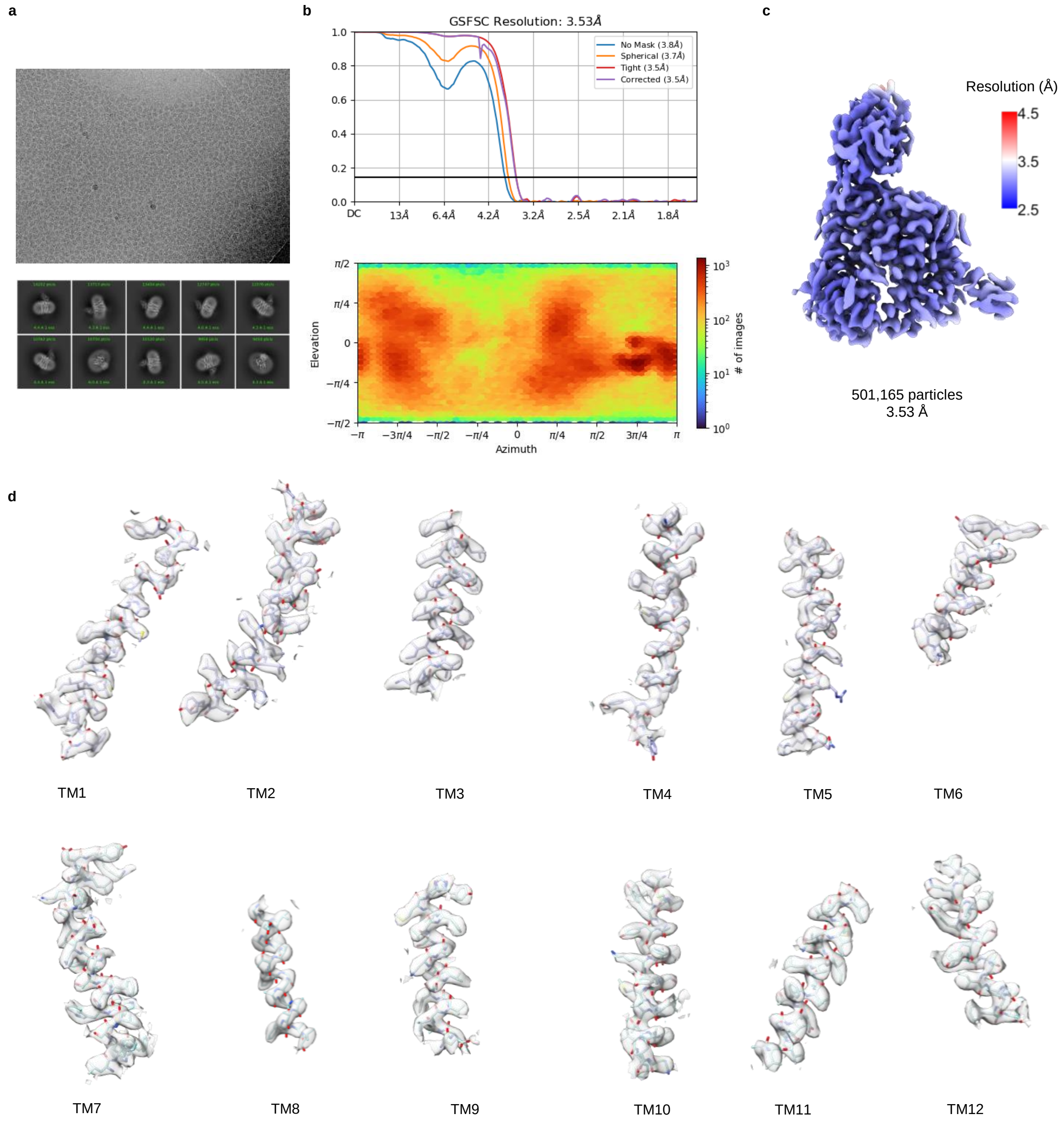


**Supplementary Fig. 9: Cryo-EM data processing overview for hSLC19A3-gf:Nb3.7:thiamine.** **a**, Representative micrograph (top) and 2D class averages (below). **b**, Gold-standard Fourier shell correlation (GSFSC) curves between two half maps (top). Angular distribution of particles in the final 3D local refinement (below). **c**, Local resolution estimation of the final cryo-EM map (map contour level = 0.151). **d**, Close-up on the density for the individual transmembrane helices (map contour level = 0.144).

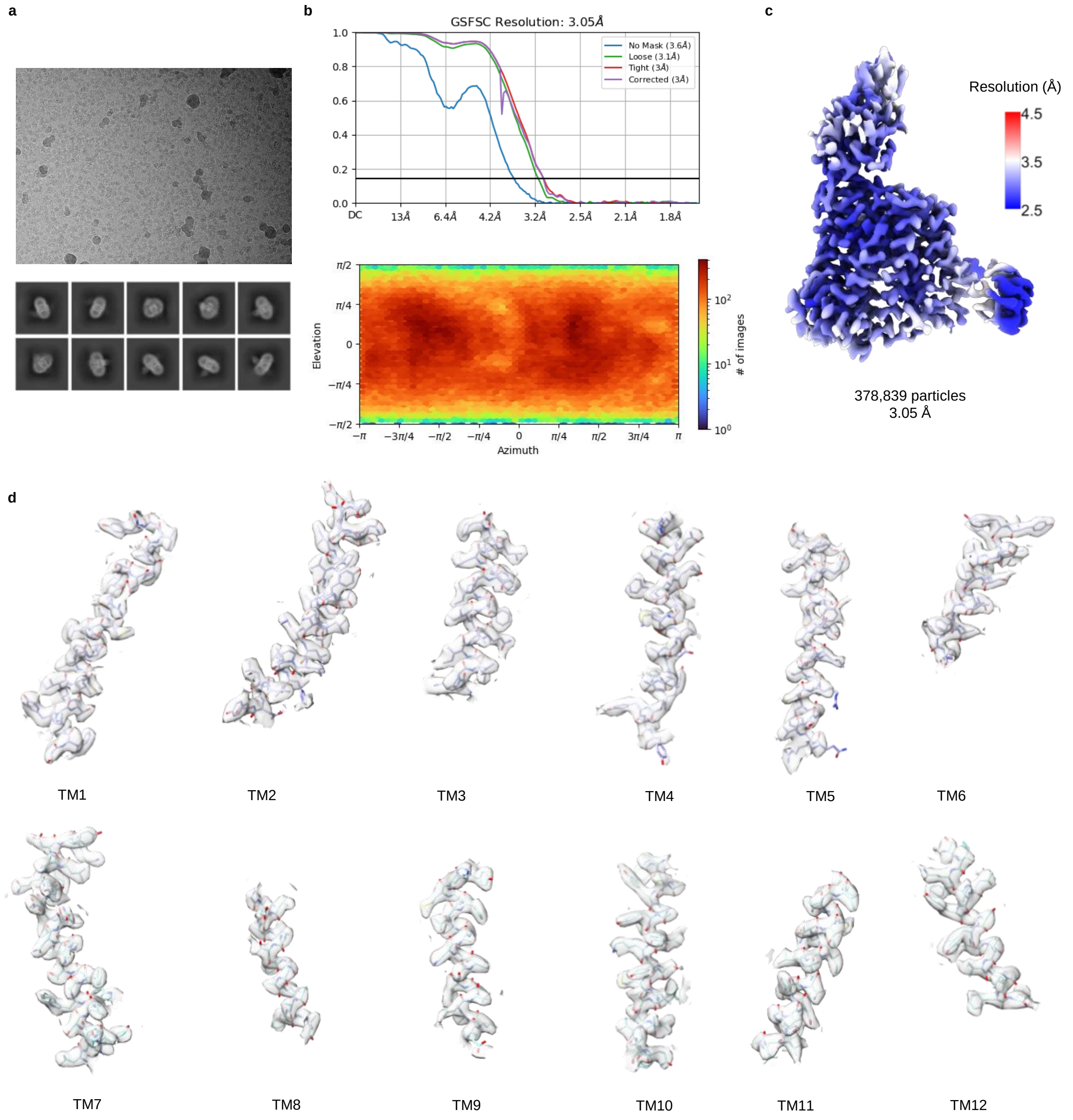


**Supplementary Fig. 10: Cryo-EM data processing overview for hSLC19A3-gf:Nb3.7:fedratinib. a**, Representative micrograph (top) and 2D class averages (below). **b**, Gold-standard Fourier shell correlation (GSFSC) curves between two half maps (top). Angular distribution of particles in the final 3D local refinement (below). **c**, Local resolution estimation of the final cryo-EM map (map contour level = 0.858). **d**, Close-up on the density for the individual transmembrane helices (map contour level = 0.563).

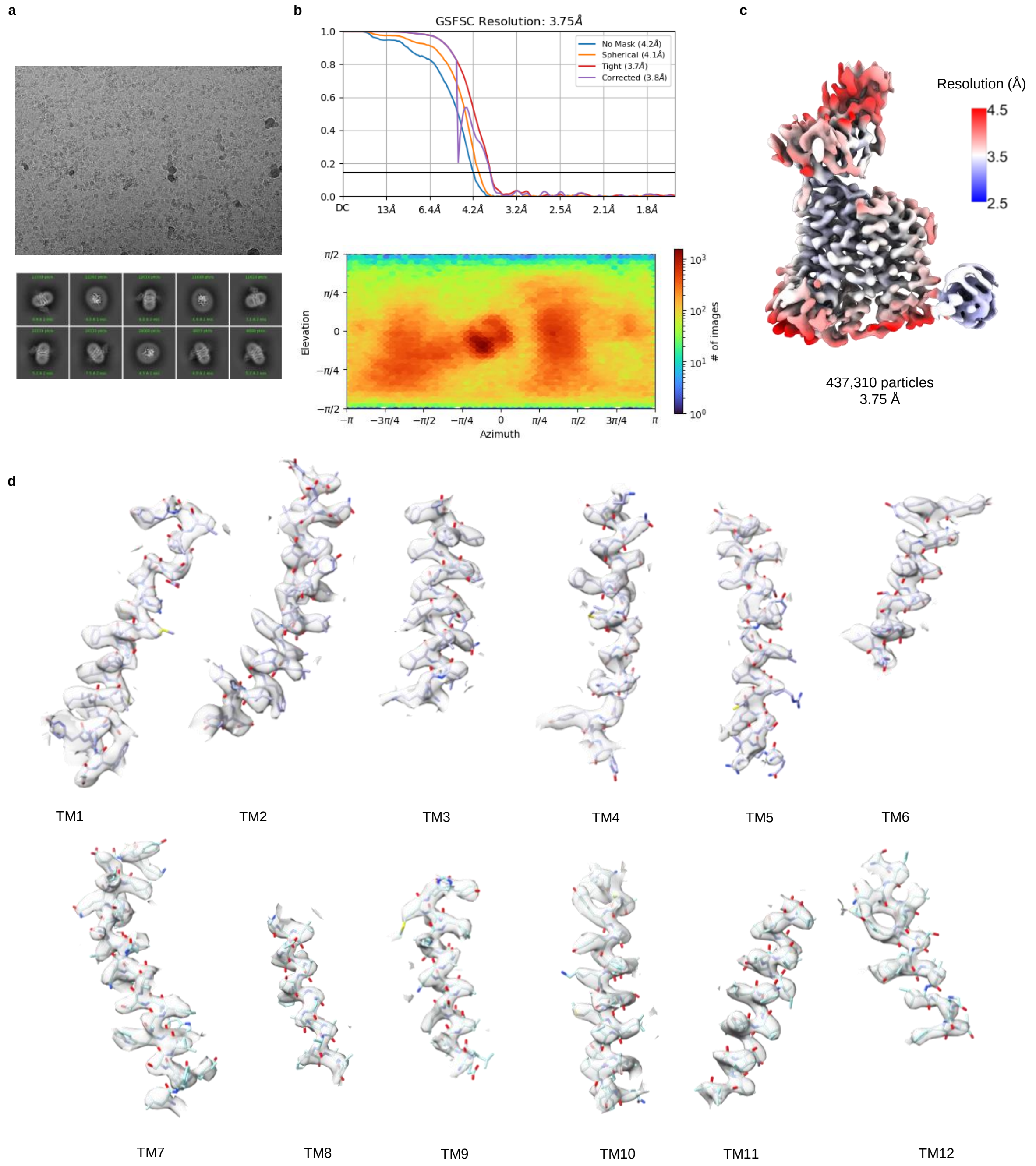


**Supplementary Fig. 11: Cryo-EM data processing overview for hSLC19A3-gf:Nb3.7:amprolium. a**, Representative micrograph (top) and 2D class averages (below). **b**, Gold-standard Fourier shell correlation (GSFSC) curves between two half maps (top). Angular distribution of particles in the final 3D local refinement (below). **c**, Local resolution estimation of the final cryo-EM map (map contour level = 0.102). **d**, Close-up on the density for the individual transmembrane helices (map contour level = 0.134).

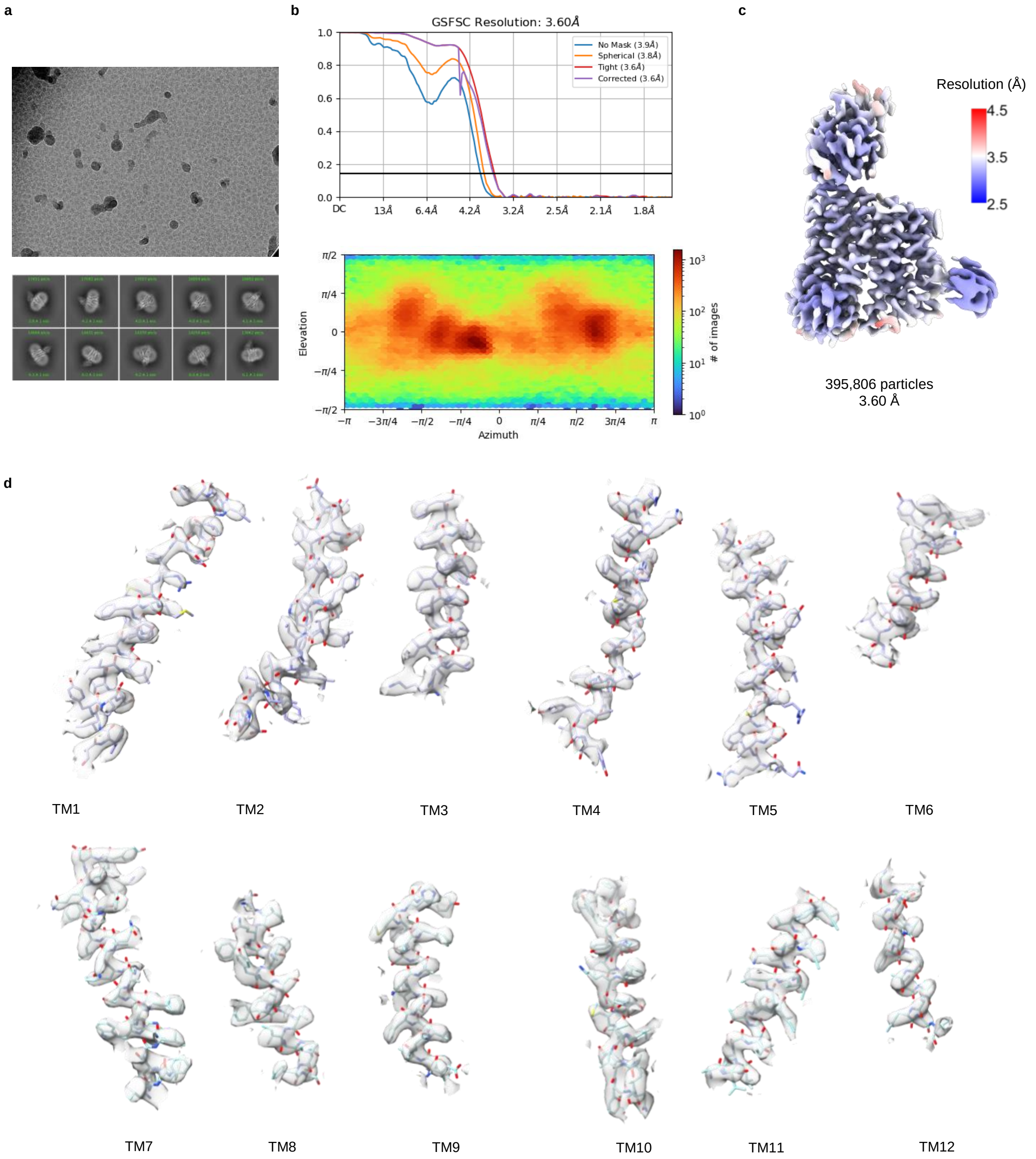


**Supplementary Fig. 12: Cryo-EM data processing overview for hSLC19A3-gf:Nb3.7:hydroxychloroquine. a**, Representative micrograph (top) and 2D class averages (below). **b**, Gold-standard Fourier shell correlation (GSFSC) curves between two half maps (top). Angular distribution of particles in the final 3D local refinement (below). **c**, Local resolution estimation of the final cryo-EM map (map contour level = 0.117). **d**, Close-up on the density for the individual transmembrane helices (map contour level = 0.133).

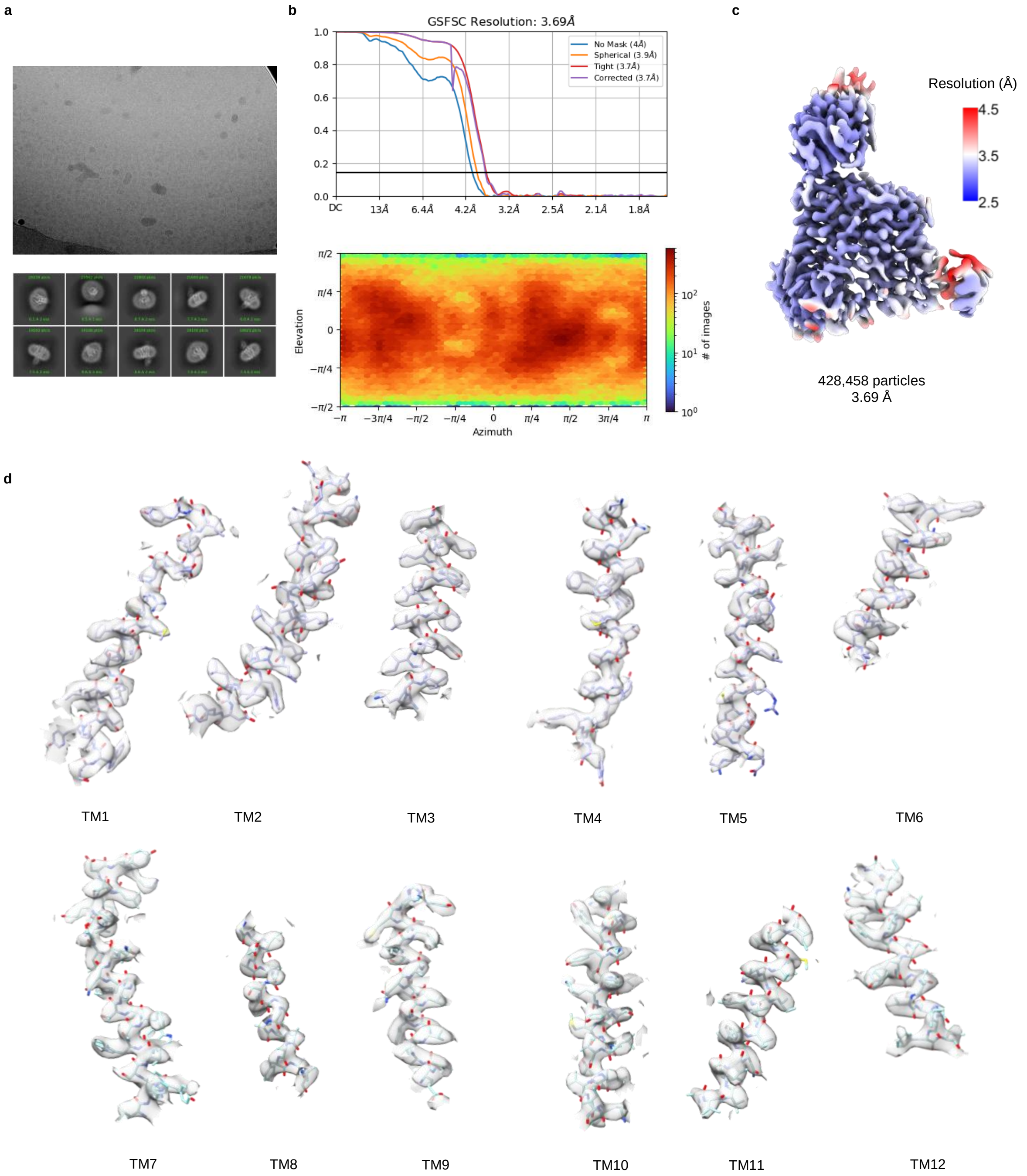


**Supplementary Fig. 13: Cryo-EM data processing overview for hSLC19A3-gf:Nb3.7:amitriptyline. a**, Representative micrograph (top) and 2D class averages (below). **b**, Gold-standard Fourier shell correlation (GSFSC) curves between two half maps (top). Angular distribution of particles in the final 3D local refinement (below). **c**, Local resolution estimation of the final cryo-EM map (map contour level = 0.856). **d**, Close-up on the density for the individual transmembrane helices (map contour level = 0.803).

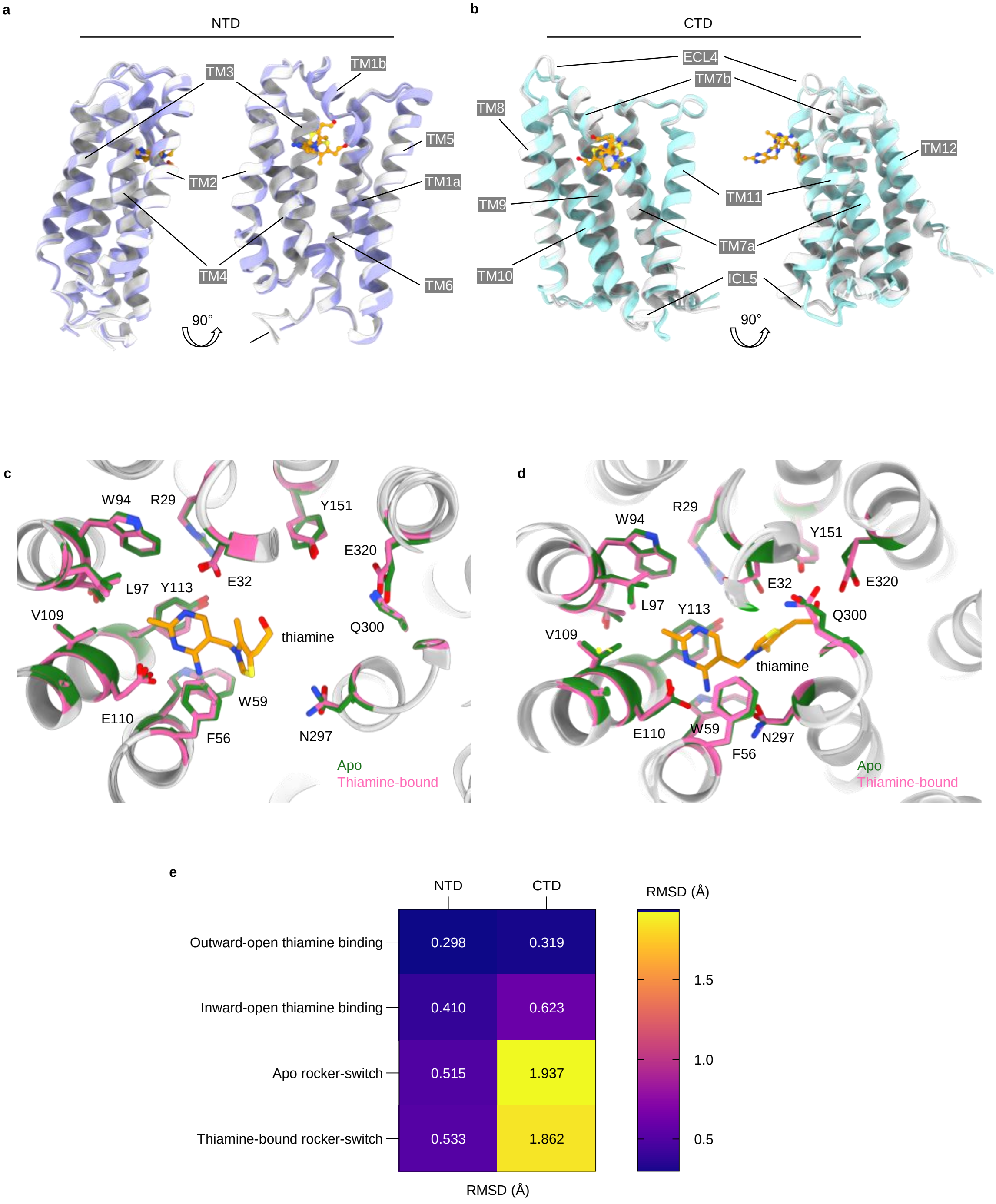


**Supplementary Fig. 14: Structural differences between different conformational states.**

**Supplementary Fig. 14: Structural differences between different conformational states. a**, **b**, Structural superposition of the protein backbones of hSLC19A3 in four different conformations, when aligned to the individual domains, **a**, NTD and **b**, CTD. The coloured structures present the outward-open states (apo and thiamine-bound). The white structures represent the inward-open states (apo and thiamine-bound). The domains move as rigid bodies between the inward- and outward-open state. The only minor intradomain changes that can be observed are rearrangements of ECL4 and ICL5. The movement of ECL4 is likely induced by the binding of Nb3.4There are no major further rearrangements observable between the apo and thiamine-bound transporter. **c** and **d** show the substrate binding site of hLSC19A3 in the outward- and inward-open state, respectively. A comparison of the apo (green) and the thiamine-bound (pink) structures reveals no major difference between them. This indicates that thiamine is recognised in a key-in-lock manner by the transporter. **e**, Assessment of the overall structural changes of the transporter between different states as root mean square deviation (RMSD) of $C_\alpha$ atoms of the individually aligned domains (NTD and CTD). Thiamine-binding results in a very low RMSD of 0.3-0.6 Å in both domains. Larger rearrangements can only be observed in the CTD, when transitioning in the rocker-switch like motion between the outward- and inward-open state. The observed higher RMSD values (1.862-1.937 Å) are mainly caused by the rearrangements in ECL4 and ICL5.

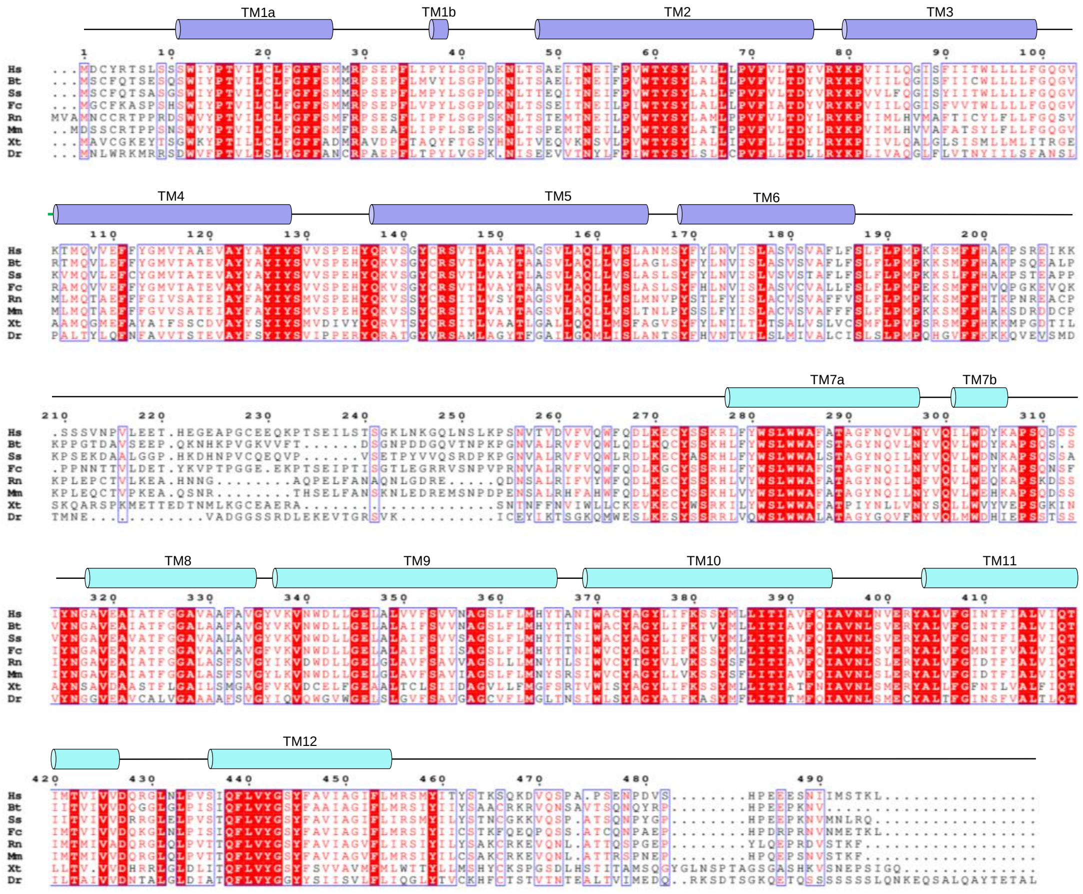

**Supplementary Fig. 15: Sequence conservation of SLC19A3 in vertebrates.** Multiple sequence alignment of SLC19A3 protein sequences of human (*Homo sapiens*, Hs), cattle (*Bos taurus*, Bt), pig (*Sus scrofa*, Ss), cat (*Felis catus*, Fs), rat (*Rattus norvegicus*, Rn), mouse (*Mus musculus*, Mm), Western clawed frog (*Xenopus tropicalis*, Xt) and zebra fish (*Danio rerio*, Dr). Residues are coloured by conservation. The topology of the transporter is indicated by the corresponding transmembrane helices (TMs) above the sequence alignment.

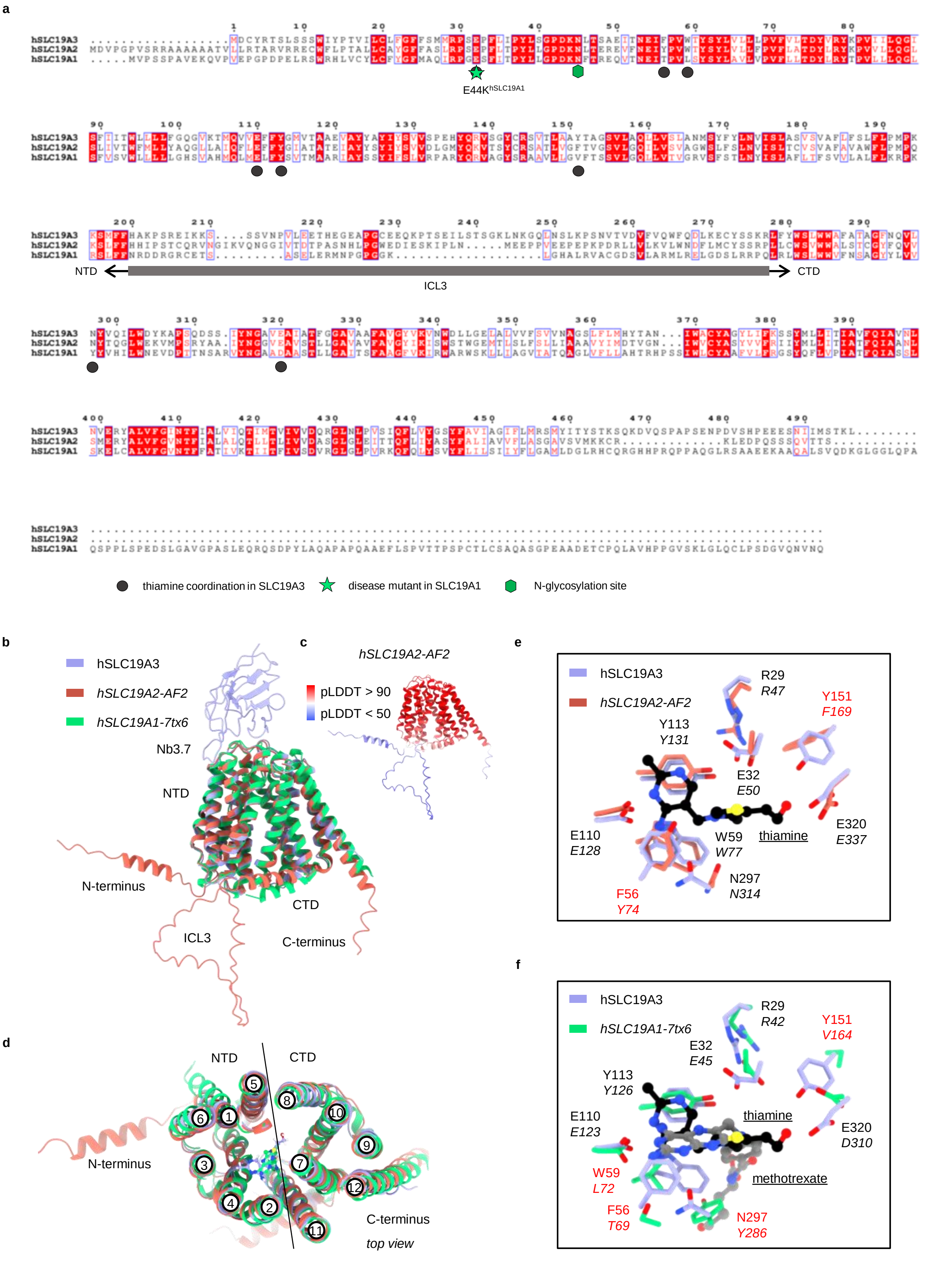


**Supplementary Fig. 16: Structure and sequence comparison between the members of the SLC19A family.**

**Supplementary Fig. 16: Structure and sequence comparison between the members of the SLC19A family. a**, Multiple sequence alignment of human SLC19A3, SLC19A2 and SLC19A1. Residue numbering is based on hSLC19A3. Residue colouring by conservation. Crucial functional or disease related residues are labelled. **b**, Side view of the superposition of the cryo-EM structures of hSLC19A3:Nb3.7:thiamine, methotrexate-bound hSLC19A1 (PDB ID 7tx6, Wright et al., 2022) and the structure model of hSLC19A2, predicted by AlphaFold2 (AF2). All structures are in the inward-open state. The fold of the three members of the SLC19A family is highly conserved. On a backbone level, only small differences can be observed. **c**, hSLC19A2 AF2 predicted structure, coloured by the predicted local distance difference test (pLDDT) score. **d**, Top view of **b**. **e**, Close-up on the substrate binding site, comparing hSLC19A3 (cryo-EM structure) and hSLC19A2 (AF2 prediction). The substrate binding site is structurally highly conserved. Only two residue substitutions can be observed from hSLC19A3 to hSLC19A2, namely Phe56 to Tyr and Tyr151 to Phe. **f**, Close-up on the substrate binding site, comparing thiamine-bound hSLC19A3 and methotrexate-bound hSLC19A1. While Glu32, Glu110 and Tyr113 are structurally conserved in hSLC19A1, the aromatic clamp is reshuffled to allow a membrane-perpendicular binding of the methotrexate aromatic ring system in the transporter. This stands in contrast to the membrane-parallel coordination of the aminopyrimidine ring of thiamine by hSLC19A3.

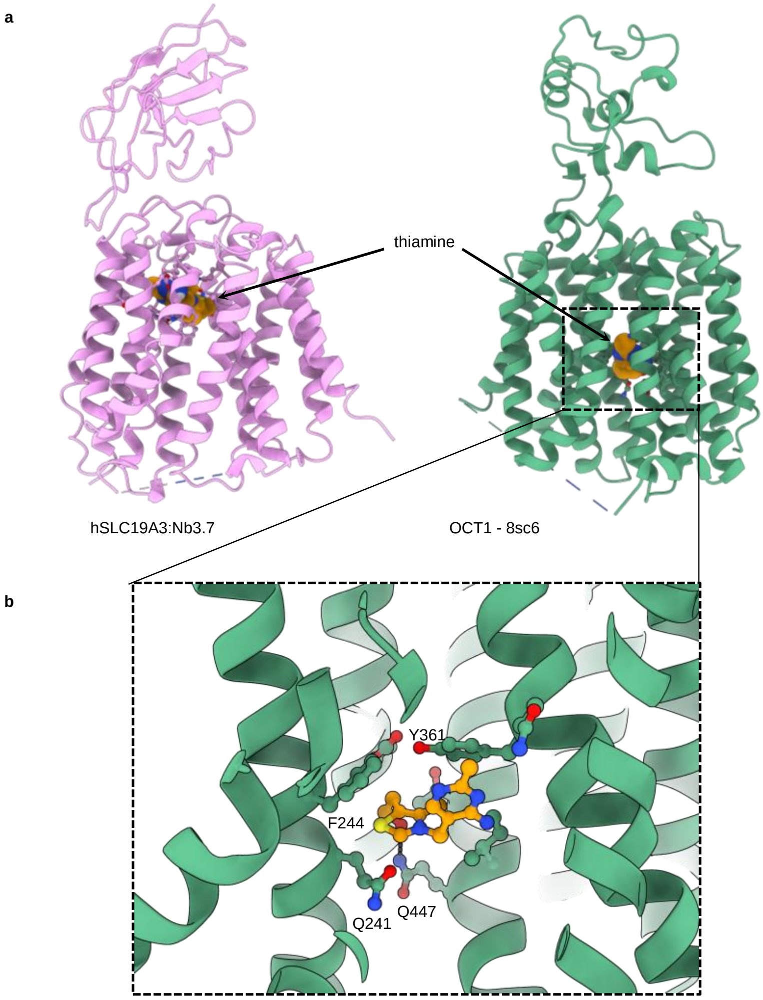


**Supplementary Fig. 17: Structure comparison between hSLC19A3 and OCT1 (hSLC22A1). a**, Side view of the cryo-EM structures of hSLC19A3:Nb3.7:thiamine and thiamine-bound OCT1 (SLC22A1, PDB ID 8sc6, Zeng et al., 2023). Thiamine is highlighted as orange sphere model. The substrate binding site of SLC19A3 is close to the extracellular space, whereas the substrate binding site of OCT1 is close to the centre of the membrane. **b**, Close-up on the substrate binding site of OCT1 coordinating thiamine. Thiamine interacts via π-π interactions with Phe244 and Tyr361 and forms a hydrogen bond to Gln447 with its hydroxyethyl tail.

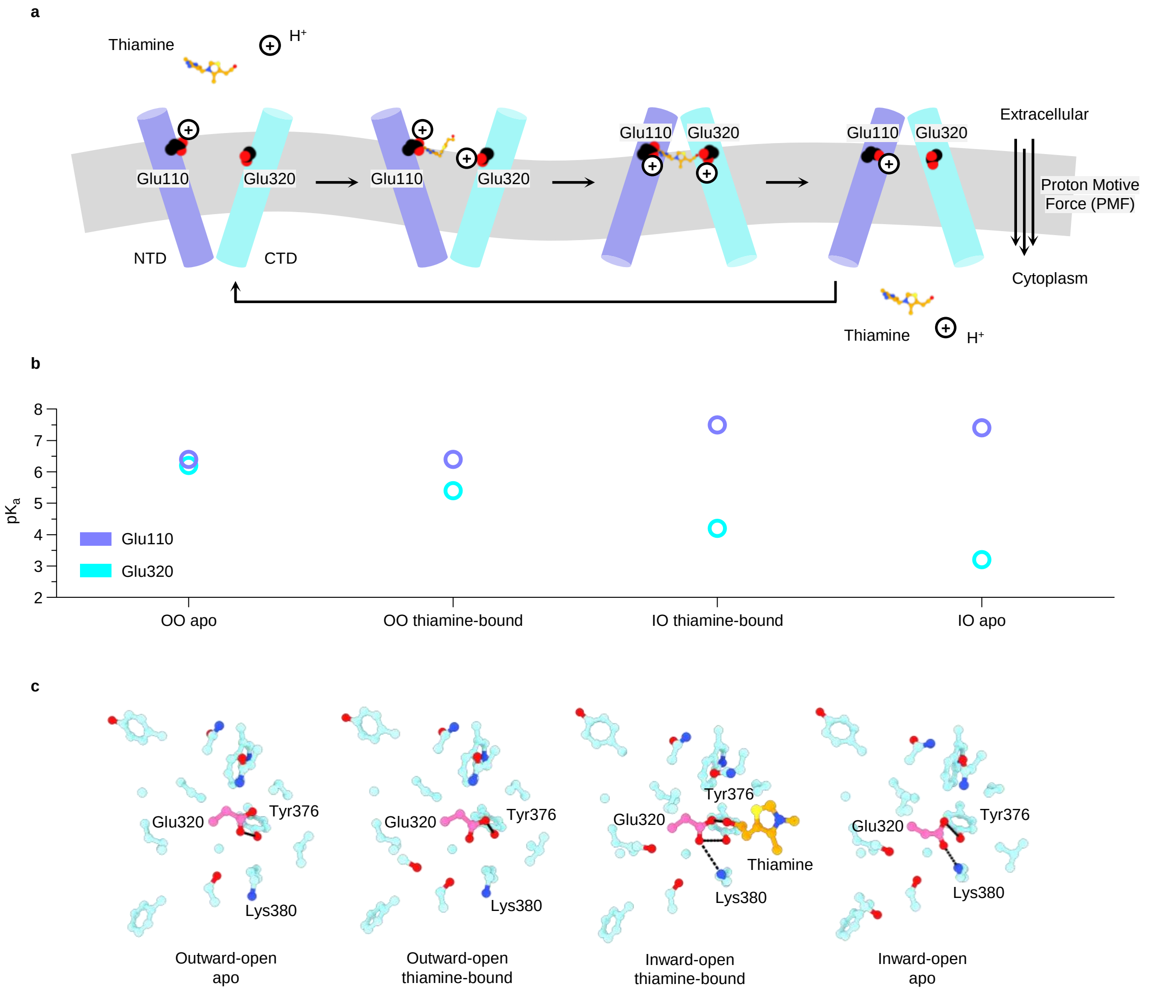


**Supplementary Fig. 18: Model for the molecular basis of proton-driven thiamine uptake. a**, Proposed protonation states of two glutamate residues critical for thiamine binding. Glu110 and Glu320, in the different conformational states of hSLC19A3 are highlighted. **b**, The proposed protonation states are based on $pK_a$ predictions using the experimentally determined structures of hSLC19A3 and the prediction software PROPKA (Bas et al., 2008). Glu110 is predicted to have a relatively constant $pK_a$ in the neutral range (6.4-7.5). Therefore, this residue is likely protonated throughout the transport cycle. Glu320, in contrast, experiences a strong $pK_a$ shift from 6.2 in the outward-open state (OO), to 3.2 in the inward-open state (IO). **c**, On a structural level, this $pK_a$ shift is likely linked to the interaction of Glu320 with Lys380. In the outward-open state, these residues are >5 Å apart and Glu320 is placed in a rather hydrophobic environment, which makes its protonation favourable. In the inward-open state, however, Glu320 and Lys380 form a salt bridge, which stabilises the ionised glutamate side chain and thus promotes the dissociation of the previously bound proton. This predicted conformation dependent protonation of Glu320 suggests that this residue could be involved in connecting thiamine uptake to the proton motive force (PMF). In the proposed model, Glu320 would bind a proton from the extracellular space in the outward-open state and release it in the cytoplasm, when it transitions to the inward-open state. The transporter could then cycle back to the outward-open state with deprotonated Glu320, thereby closing the co-transport cycle of a proton. In this model, co-transport of a proton would be an additional driving force for thiamine uptake.

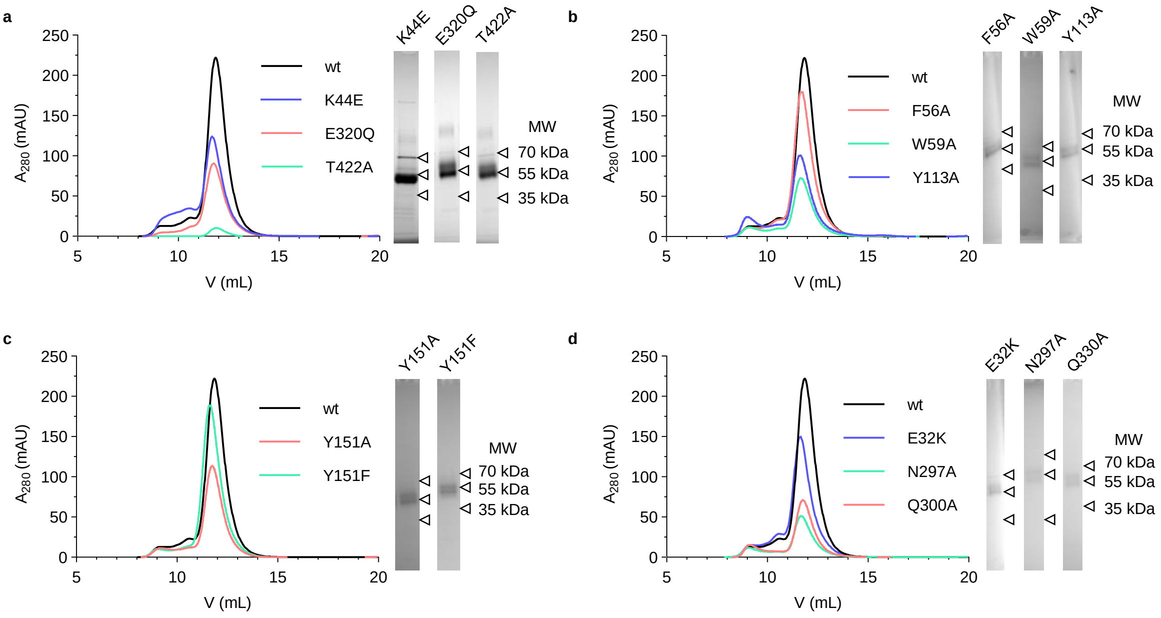


**Supplementary Fig. 19: Purification of single point mutants of hSLC19A3. a-d**, Size-exclusion chromatography (SEC) traces are shown in the respective chromatograms, while the insets show SDS-PAGE analysis of the pooled SEC-peak fraction. For **a**, concentrated protein was loaded on the SDS-PAGE gels, whereas for the rest of the shown gels, the unconcentrated peak fraction was loaded directly. Note in **a**, left inset, that the disease-causing mutant Lys44Glu lacks the additional higher molecular weight band slightly above 55 kDa, which represents glycosylated protein. It has been shown previously, that this mutant is retained in the endoplasmic reticulum (ER, Kono et al., 2009). Our data exhibit that hSLC19A3-Lys44Glu is likely not properly glycosylated in the ER which could prevent trafficking to the plasma membrane. For further details, see Supplementary Fig. 23b. White triangles indicate the molecular weight markers.

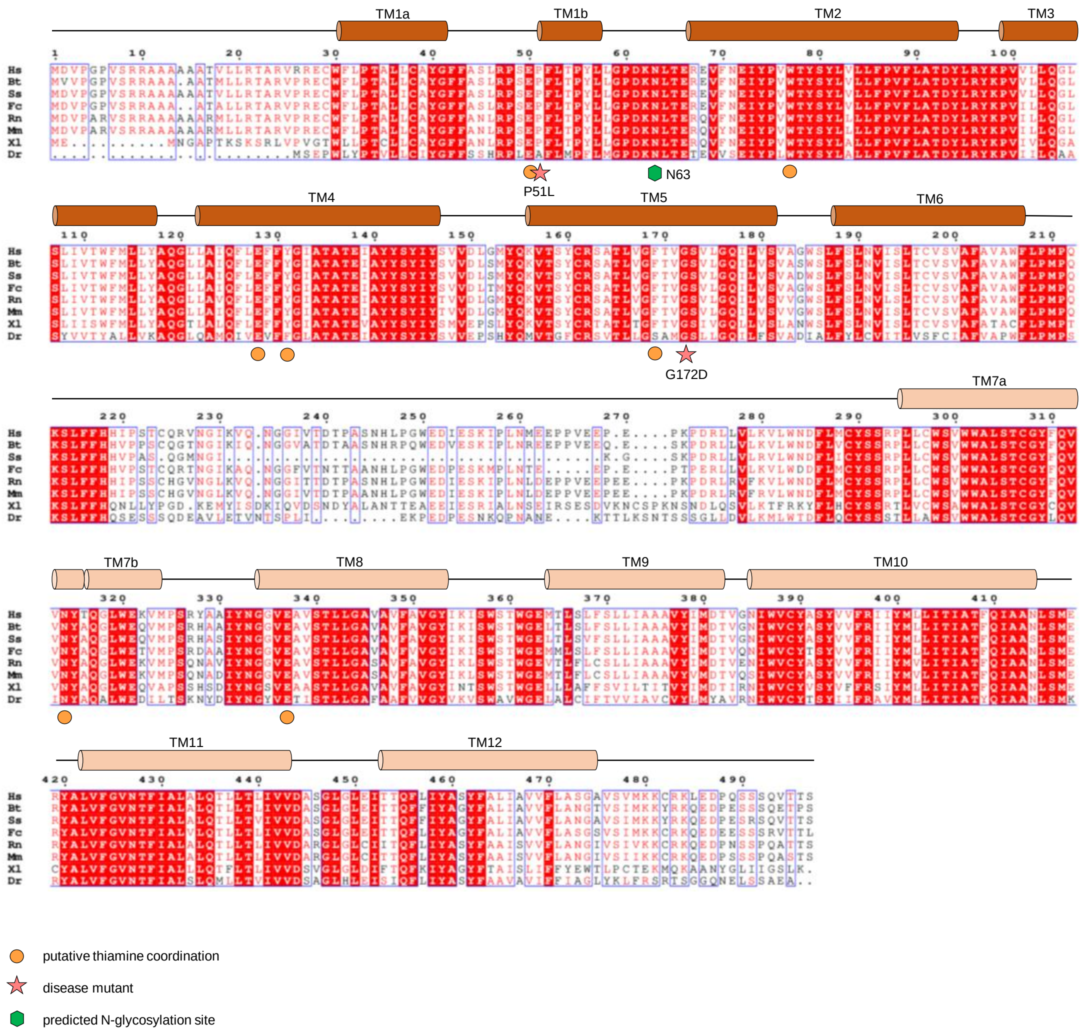


**Supplementary Fig. 20: Sequence conservation of SLC19A2 in vertebrates.** Multiple sequence alignment of SLC19A2 protein sequences of human (*Homo sapiens*, Hs), cattle (*Bos taurus*, Bt), pig (*Sus scrofa*, Ss), cat (*Felis catus*, Fs), rat (*Rattus norvegicus*, Rn), mouse (*Mus musculus*, Mm), Western clawed frog (*Xenopus tropicalis*, Xt) and zebra fish (*Danio rerio*, Dr). Residues are coloured by conservation. The topology of the transporter is indicated by the corresponding transmembrane helices (TMs) above the sequence alignment. The stars indicate mutants of hSLC19A2, which cause thiamine-responsive megaloblastic anemia (TRMA), namely Pro51Leu (Lagard et al., 2004) and Gly172Asp (Labay et al., 1999). Residues likely important for substrate coordination (based on homology to SLC19A3) are labelled with an orange circle.

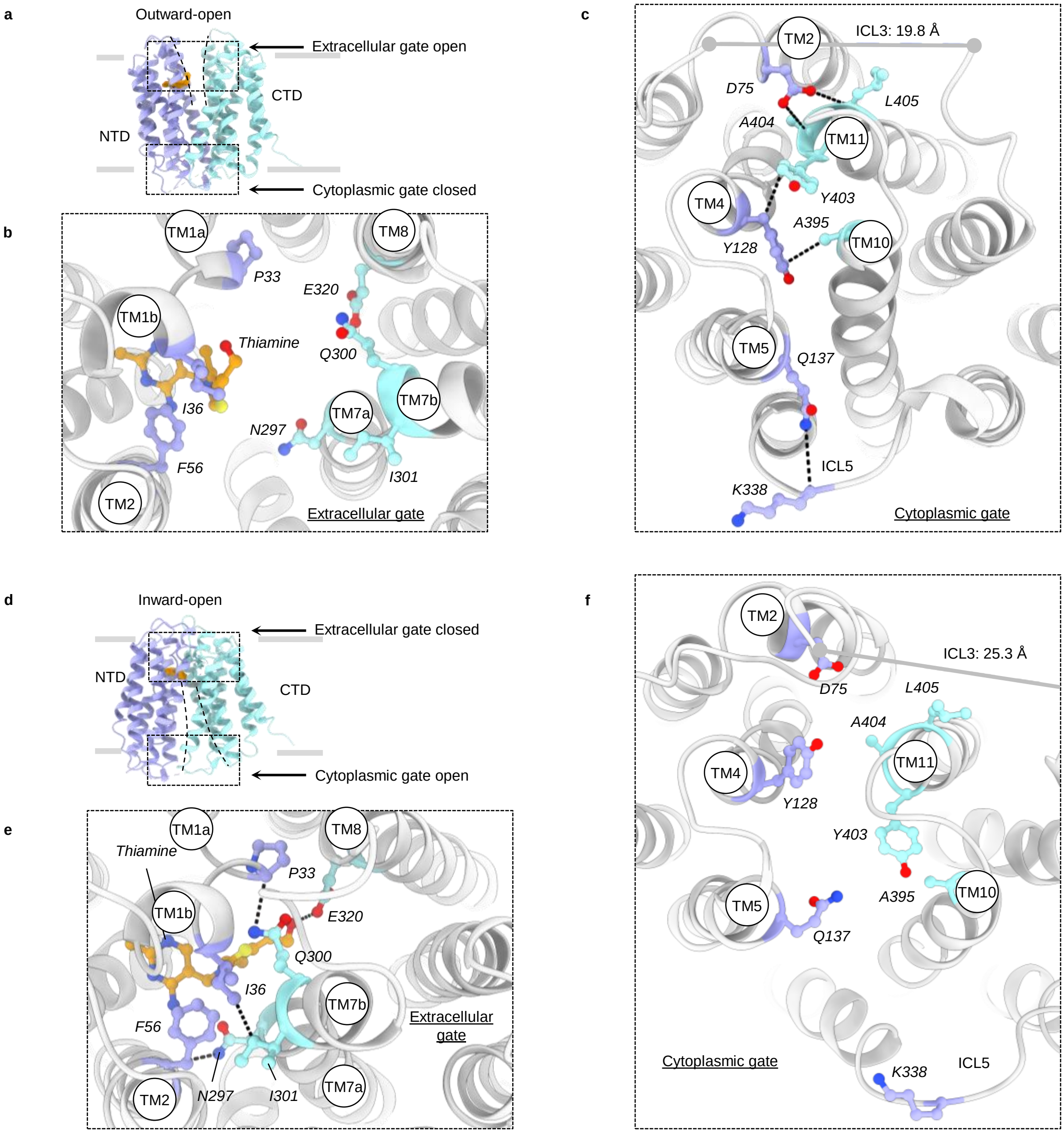


**Supplementary Fig. 21: Molecular gates. a-c**, Representation of the molecular gates in the outward-open state of hSLC19A3. While the extracellular gate is open, the cytoplasmic gate is sealed by the apolar interactions between the side chains of Tyr128 on the NTD and Ala395 and Tyr403 on the CTD. The gate is further stabilised by polar interactions between Asp75 and the backbone of Leu405, as well as Gln137 and the backbone of Lys338. **d-f**, Molecular gates of hSLC19A3 in the inward-open state. The extracellular gate is closed. Central polar contacts are formed by Asn297 with the backbone of Phe56 and Gln300 and the backbone of Pro33. The gate is further sealed by the apolar interaction of Ile36 with Ile301. The cytoplasmic gate is open. The structured endpoints of ICL3 move away from each other (from ~20 Å to about 25 Å, measured between $C_\alpha$ of 195 and 271). Selected residues are shown as sticks and black dashes indicate hydrogen bonds and hydrophobic interactions (cut-off at 3.2 Å)

**a**

| Disease | Age of onset | Mutation | Structural module | Specific reference |
|---|---|---|---|---|
| **Classical childhood BTBGD** | 4.73 ± 0.35 years | P15L | | |
| | | G41R | | |
| | | W59R | Substrate binding site | Aburezq et al., 2023 |
| | | L66P | | |
| | | W94R | Substrate binding site | Schänzer et al., 2014 |
| | | G100R | | |
| | | Y113H | Substrate binding site | |
| | | V139E | | |
| | | S155L | | |
| | | S163P | | |
| | | N173D | | |
| | | S176Y | | Kevelam et al., 2013 |
| | | S181P | | Kevelam et al., 2013 |
| | | W284R | | |
| | | G317E | | |
| | | E320K | Substrate binding site | Wesol-Kucharska et al., 2021 |
| | | A321V | | |
| | | L385R | | Kevelam et al., 2013 |
| | | N399I | | |
| | | V400L | | |
| | | T422A | | |
| **Infantile phenotype** | 3.00 ± 0.48 months | P33Q | EC gate | |
| | | L46P | | |
| | | E120K | | |
| | | A183V | | |
| | | I301V | EC gate | |
| | | N316T | | |
| | | L385R | | |
| | | S444R | | |
| **Mixed type** | | G23V | | |
| | | S89R | | |
| **WLE** | 10-20 years | K44E | | Kono et al., 2009 |
| | | E320Q | Substrate binding site | Kono et al., 2009 |

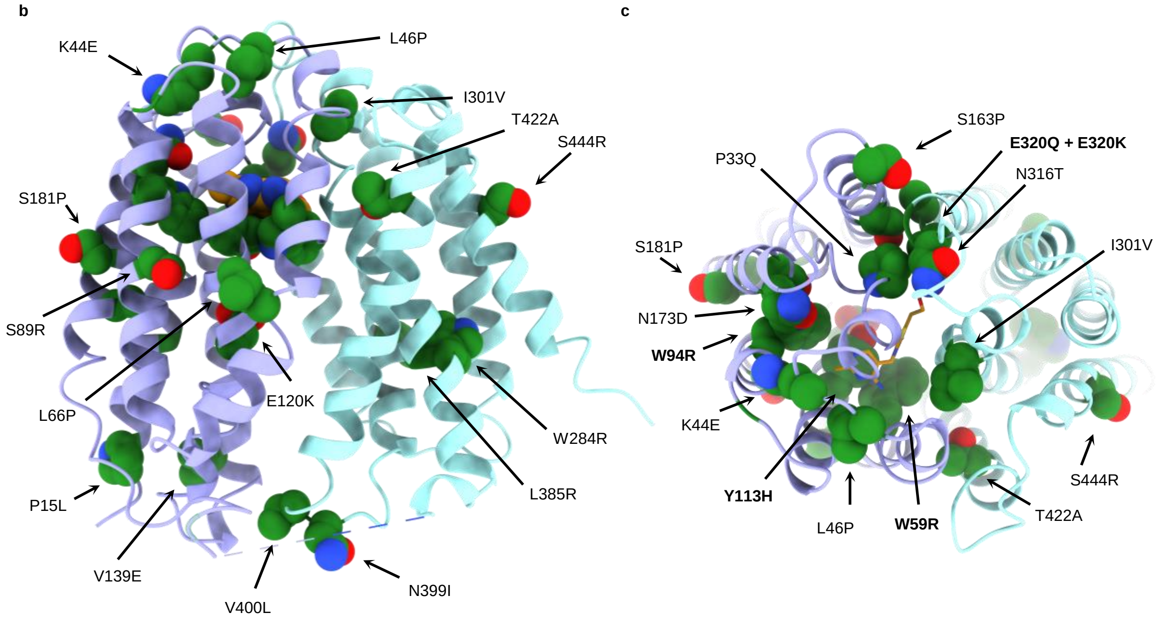


**Supplementary Fig. 22: Disease-causing mutations of hSLC19A3.**

**Supplementary Fig. 22: Disease-causing mutations of hSLC19A3. a**, List of disease-causing point mutants of hSLC19A3. Most of these mutants cause biotin- and thiamine-responsive basal ganglia disease (BTBGD). This disease can be subdivided in classical childhood BTBGD, BTBGD with infantile phenotype and mixed type BTBGD, based on the age of onset (Wang et al., 2021). The heterozygous combination of the two latter mutations (K44E and E320Q) causes in contrast Wernicke's like encephalopathy (WLE), with an onset of symptoms in the second decade of life (Kono et al., 2009). Unless otherwise specified, see Want et al., 2021 for a review of the mutations. Based on our structural data, we can assign a structural function to several of the mutated residues, e.g. when they are part of the substrate binding site or the molecular gates of the transporter. **b**, Side view of the cryo-EM structure of inward-open hSLC19A3 bound to thiamine (orange). Disease causing mutations as listed in the table above are highlighted in dark green sphere in the structure. **c**, Top view of the same complex. Substrate binding site residues are highlighted in bold font.

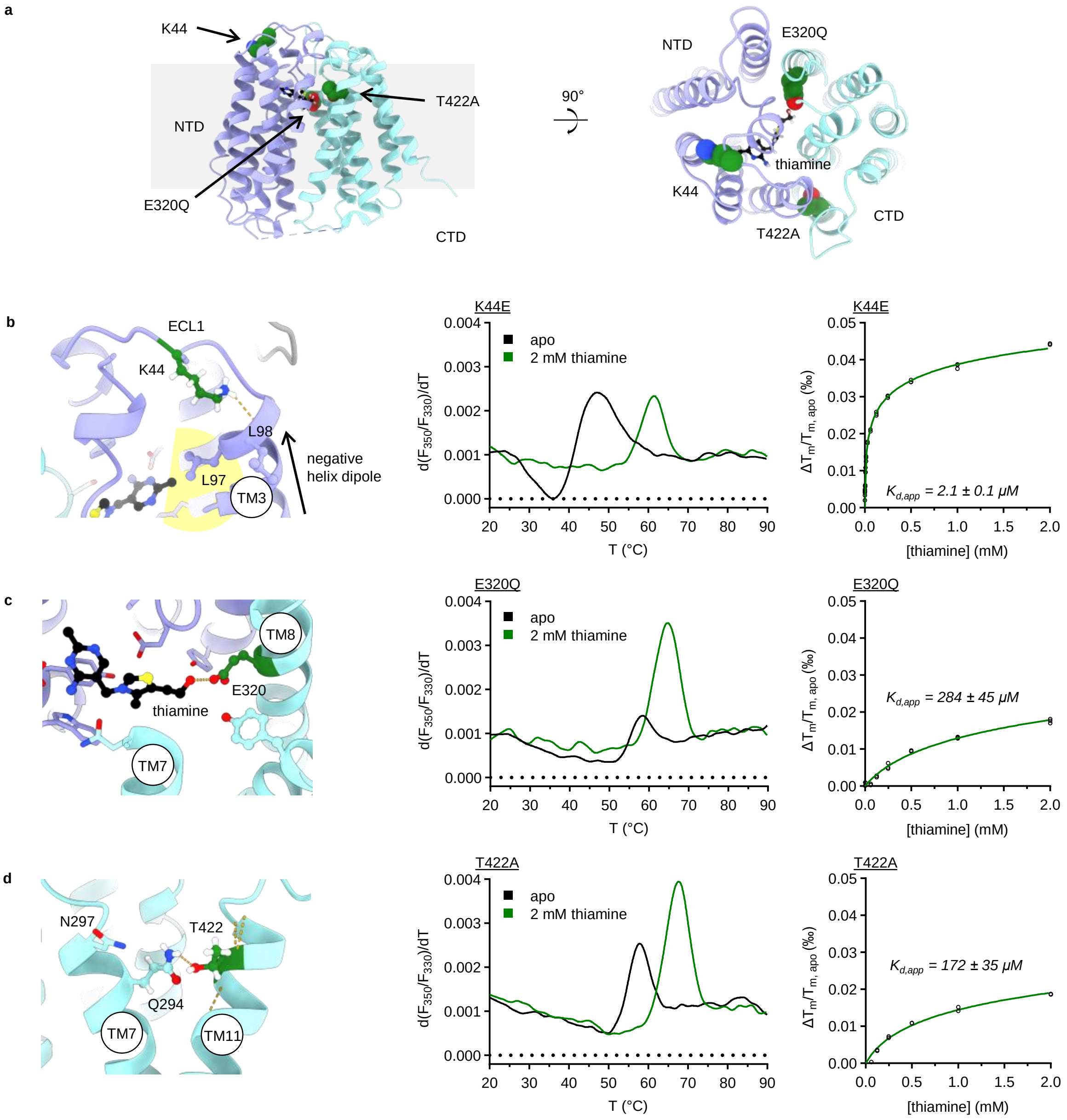


**Supplementary Fig. 23: Thermostability and thiamine binding affinity of selected disease-causing mutations of hSLC19A3.**

**Supplementary Fig. 23: Thermostability and thiamine binding affinity of selected disease-causing mutations of hSLC19A3. a**, Localisation of the studied disease-causing mutations. Individuals that carry aSLC19A3-Lys44Glu and -Glu320Gln allele at the same time develop Wernicke's like encephalopathy (Kono et al., 2009). Thr422Ala is the most reported single point mutation of SLC19A3 causing biotin- and thiamine-responsive basal ganglia disease (BTBGD) in homozygous individuals. **b-d**, The panels below show structural close-ups to the affected residues in the cryo-EM structure of hSLC19A3:Nb3.7:thiamine (left), representative melting curves (middle; first derivative of the ratio of the fluorescence at 350 nm and 330 nm (F350/F330)), and thiamine concentration dependent thermal shifts used for the determination of thiamine binding affinities (right). **b**, Lys44 forms polar contacts with the C-terminal end of TM3. Its positively charged side chain engages with the backbone carbonyl of Leu98 and presumably with the negative helix dipole of TM3. The thermostability of this mutant is strongly reduced ($\Delta T_m^{K44E\text{-}wt}$ = -15 °C), when compared to wildtype. However, the mutated transporter binds thiamine with a six-fold higher affinity than wildtype hSLC19A3. While this mutant appears to retain its substrate binding capacity, it has been shown that it cannot be trafficked properly to the plasma membrane (Kono et al., 2009). Our data, in the form of a reduced thermostability and lacking glycosylation of hSLC19A3-Lys44Glu (Supplementary Fig. 19a) suggest that the mutant might be quite unstable to pass protein quality control in the endoplasmic reticulum (ER). **c**, hSLC19A3-Glu320Gln, in contrast, localises to the plasma membrane (Kono et al., 2009); the thiamine uptake via this mutant is, however, strongly impaired. Our biophysical data display, that the stability of the mutant is only moderately affected ($\Delta T_m^{E320Q\text{-}wt}$ = -4 °C). The affinity for thiamine was, however, markedly reduced from a $K_{d,app}$ of 12.7 μM to 284 μM. This underpins the importance of the hydrogen bond between Glu320 and the hydroxyethyl tail of thiamine for high-affinity binding. **d**, Thr422 forms a hydrogen bond with Gln294, which might be important for proper helix packing. The thermostability of the Thr422Ala mutant of hSLC19A3 is decreased and binds thiamine with > ten-fold lower affinity. The lower binding affinity could render the transporter unreceptive for physiological thiamine levels. It might, however, be able to transport thiamine at higher concentrations, which is supported by the success of high-dose thiamine treatment of patients carrying this mutation.

**Supplementary Table 1, part 1 of 2 | Cryo-EM data collection, refinement and validation statistics**

| Structure<br>PDB<br>EMDB | hSLC19A3-wt:Nb3.4-apo<br>8S4U<br>EMD-19716 | hSLC19A3-wt:Nb3.4:thiamine<br>8S5U<br>EMD-19750 | hSLC19A3-wt:Nb3.3-apo<br>8S5V<br>EMD-19751 | hSLC19A3-gf:Nb3.7:thiamine<br>8S61<br>EMD-19754 |
|---|---|---|---|---|
| **Data collection/ processing** | | | | |
| Magnification | 105,000x | 105,000x | 105,000x | 105,000x |
| Voltage (kV) | 300 | 300 | 300 | 300 |
| Pixel size (Å) | 0.83 | 0.85 | 0.83 | 0.83 |
| Defocus range (μM) | 0.8-1.6 | 0.8-1.6 | 0.8-1.6 | 0.8-1.6 |
| Electron exposure ($e^-$/Å$^2$) | 45 | 62 | 45 | 45 |
| Symmetry imposed | C1 | C1 | C1 | C1 |
| Movies (No.) | 5,859 | 9,105 | 6,508 | 5,308 |
| Initial particles (No.) | 1,084,118 | 1,602,998 | 1,619,839 | 1,264,486 |
| Final particles (No.) | 486,079 | 277,942 | 512,258 | 501,165 |
| Map resolution (Å) | 3.09 | 3.28 | 2.87 | 3.53 |
| FSC threshold | 0.143 | 0.143 | 0.143 | 0.143 |
| Map resolution range (Å) | 2.6-3.5 | 2.8-4.5 | 2.4-3.0 | 2.9-3.4 |
| **Refinement** | | | | |
| Initial model used (PDB code) | AF_AFQ9BZV2F1 | AF_AFQ9BZV2F1 | AF_AFQ9BZV2F1 | AF_AFQ9BZV2F1 |
| Model resolution (Å) | 3.1 | 3.3 | 3.0 | 3.6 |
| FSC threshold | 0.5 | 0.5 | 0.5 | 0.5 |
| Map sharpening B-factor (Å$^2$) | -65.6 | -90.1 | -80.0 | -111.1 |
| **Model composition** | | | | |
| Non-hydrogen atoms | 3,981 | 3,999 | 4,042 | 3,990 |
| Protein residues | 502 | 502 | 507 | 502 |
| Ligands | 1 | 2 | 1 | 1 |
| **B factors (Å$^2$)** | | | | |
| Protein | 29.14 | 27.76 | 27.31 | 10.58 |
| Ligand | 30.00 | 24.38 | 30.00 | 10.30 |
| **R.m.s. deviations** | | | | |
| Bond lengths (Å) | 0.006 | 0.005 | 0.004 | 0.004 |
| Bond angles (°) | 0.627 | 0.576 | 0.617 | 0.540 |
| **Validation** | | | | |
| MolProbity score | 1.72 | 1.57 | 1.29 | 1.72 |
| Clashscore | 7.41 | 5.64 | 4.22 | 5.41 |
| Rotamer outliers (%) | 0.24 | 0.47 | 0.94 | 0.00 |
| **Ramachandran plot (%)** | | | | |
| Favoured | 95.56 | 96.17 | 97.60 | 93.35 |
| Allowed | 4.44 | 3.83 | 2.20 | 6.65 |
| Outliers | 0.00 | 0.00 | 0.20 | 0.00 |

**Supplementary Table 1, part 2 of 2 | Cryo-EM data collection, refinement and validation statistics**

| Structure<br><br>PDB<br><br>EMDB | hSLC19A3-gf:Nb3.7: Fedratinib<br><br>8S5W<br><br>EMD-19752 | hSLC19A3-gf:Nb3.7: Amprolium<br><br>8S62<br><br>EMD-19755 | hSLC19A3-gf:Nb3.7: Hydroxy-chloroquine<br><br>8S5Z<br><br>EMD-19753 | hSLC19A3-gf:Nb3.7: Amitriptyline<br><br>8S66<br><br>EMD-19756 |
|---|---|---|---|---|
| **Data collection/ processing** | | | | |
| Magnification | 105,000x | 105,000x | 105,000x | 105,000x |
| Voltage (kV) | 300 | 300 | 300 | 300 |
| Pixel size (Å) | 0.83 | 0.83 | 0.83 | 0.83 |
| Defocus range (μM) | 0.8-1.6 | 1.0-2.0 | 0.8-1.6 | 0.8-1.6 |
| Electron exposure ($e^-$/Å$^2$) | 45 | 46 | 52 | 51 |
| Symmetry imposed | C1 | C1 | C1 | C1 |
| Movies (No.) | 5,328 | 5,794 | 5,047 | 15,588 |
| Initial particles (No.) | 1,303,193 | 932,296 | 1,334,920 | 1,921,555 |
| Final particles (No.) | 378,839 | 437,310 | 395,806 | 428,458 |
| Map resolution (Å) | 3.05 | 3.75 | 3.60 | 3.69 |
| FSC threshold | 0.143 | 0.143 | 0.143 | 0.143 |
| Map resolution range (Å) | 2.5-3.5 | 3.2-4.5 | 3.0-4.0 | 3.0-4.0 |
| **Refinement** | | | | |
| Initial model used (PDB code) | AF_AFQ9BZV2F1 | AF_AFQ9BZV2F1 | AF_AFQ9BZV2F1 | AF_AFQ9BZV2F1 |
| Model resolution (Å) | 3.1 | 4.1 | 3.9 | 3.9 |
| FSC threshold | 0.5 | 0.5 | 0.5 | 0.5 |
| Map sharpening B-factor (Å$^2$) | -80.1 | -190.9 | -127.5 | -128.0 |
| **Model composition** | | | | |
| Non-hydrogen atoms | 4,009 | 3,990 | 3,995 | 3,993 |
| Protein residues | 502 | 502 | 502 | 502 |
| Ligands | 1 | 1 | 1 | 1 |
| **B factors (Å$^2$)** | | | | |
| Protein | 30.02 | 58.38 | 19.10 | 21.62 |
| Ligand | 33.10 | 51.76 | 21.38 | 24.05 |
| **R.m.s. deviations** | | | | |
| Bond lengths (Å) | 0.006 | 0.021 | 0.007 | 0.006 |
| Bond angles (°) | 0.705 | 1.496 | 0.811 | 0.848 |
| **Validation** | | | | |
| MolProbity score | 1.71 | 1.88 | 1.94 | 1.88 |
| Clashscore | 5.26 | 9.06 | 7.79 | 7.92 |
| Rotamer outliers (%) | 0.24 | 0.24 | 0.24 | 0.48 |
| **Ramachandran plot (%)** | | | | |
| Favoured | 93.55 | 94.15 | 91.33 | 92.94 |
| Allowed | 6.45 | 5.85 | 8.67 | 6.85 |
| Outliers | 0.00 | 0.00 | 0.00 | 0.20 |
